# Tracking the mind’s eye: Primate gaze behavior during virtual visuomotor navigation reflects belief dynamics

**DOI:** 10.1101/689786

**Authors:** Kaushik J Lakshminarasimhan, Eric Avila, Erin Neyhart, Gregory C DeAngelis, Xaq Pitkow, Dora E Angelaki

## Abstract

To take the best actions, we often need to maintain and update beliefs about variables that cannot be directly observed. To understand the principles underlying such belief updates, we need tools to uncover subjects’ belief dynamics from natural behaviour. We tested whether eye movements could be used to infer subjects’ beliefs about latent variables using a naturalistic, visuomotor navigation task. We observed eye movements that appeared to continuously track the goal location even when no visible target was present there. Accurate goal-tracking was associated with improved task performance, and inhibiting eye movements in humans impaired navigation precision. By using passive stimulus playback and manipulating stimulus reliability, we show that subjects’ eye movements are likely voluntary, rather than reflexive. These results suggest that gaze dynamics play a key role in action-selection during challenging visuomotor behaviours, and may possibly serve as a window into the subject’s dynamically evolving internal beliefs.

## INTRODUCTION

Rational behaviour in the real world often requires predicting latent states from sensory observations. Since latent variables cannot be directly observed, and since the utility of actions depends on the status of latent variables in the future, subjects must use statistical regularities in space and in time to predict them. There is a growing body of studies that not only demonstrate that humans exploit regularities in feature space (Langer and Bülthoff, 2001; Miyazaki, 2005; Weiss et al., 2002), but also show how to infer the associated subjective priors from data (Gosselin and Schyns, 2003; Houlsby et al., 2013; Körding and Wolpert, 2004; Paninski, 2006; Smith et al., 2012; Stocker and Simoncelli, 2006; Turnham et al., 2011). In contrast, we know relatively little about how physical laws that govern the temporal dynamics of inputs are internalized and used to guide time-evolving beliefs in the absence of reliable observations (Lee et al., 2014).

The reasons for limited progress in understanding belief dynamics are twofold. First, psychophysics continues to be dominated by experimental paradigms in which actions are discrete (*e.g.*, binary choice) and sporadic (*e.g.*, at the end of the trial). In contrast, continuous tasks (Bonnen et al., 2015; Huk et al., 2018; Knöll et al., 2018; Pitkow and Angelaki, 2017) provide subjects the opportunity to reveal more information about their beliefs and predictions as they unfold in time. Second, although theoretical techniques to infer latent beliefs from actions are slowly becoming available (Kumar et al., 2019; Reddy et al., 2018; Wu et al., 2018, 2019), they have yet to be successfully applied to settings in which state and action spaces are both continuous. Consequently, principled ways to reliably uncover subjects’ belief dynamics from natural behaviour are still lacking. Meanwhile, a practical way to overcome this hurdle would be by covertly ‘measuring’ those beliefs. One candidate tool to accomplish this is eye-tracking (Spivey, 2007). Saccadic eye movements have previously been used to understand mental processes underlying a wide variety of abstract tasks such as language comprehension (Tanenhaus et al., 1995), reading (Rayner, 1998), mental imagery (Spivey and Geng, 2001), visual search (Zhang et al., 2018), and even random number generation (Loetscher et al., 2010). Furthermore, it has recently been argued that smooth-pursuit eye movements may be influenced by short-term memory (Deravet et al., 2018; Orban de Xivry et al., 2013). By formulating oculomotor pursuit to transiently occluded moving targets as an active inference process, these eye movements have been used to infer subjects’ internal beliefs (Adams et al., 2012, 2015). We wanted to know whether eye movements might also reflect belief dynamics for extended periods of time under more naturalistic conditions.

To address this, we first designed a challenging, naturalistic visuomotor task. We created a virtual environment comprised solely of sparse optic flow cues in which subjects used a joystick to steer to a transiently cued target location by integrating optic flow. To successfully perform the task, subjects had to continuously update an internal estimate of the relative target location by inferring their own movements based on the sparse cues. Although simply extinguishing the target does not make the target location a latent variable, the location of the target *relative* to the subject becomes latent as soon as they start steering. This is because thereafter, the relative location is not directly observed, only inferred by discounting one’s own displacements. To test whether eye movements were informative about those inferred estimates, we recorded the gaze behaviour of humans and rhesus macaques while they performed this task. Parallel experiments in the two species allowed us to test whether the observed eye movements were evolutionarily conserved. We found that both humans and monkeys tend to follow the location of the unseen target with their gaze until they reach it. By manipulating stimulus reliability and by using stimulus playback, we demonstrate that the eye movements are likely volitional, rather than reflexive. Furthermore, the subjects’ success in tracking the target over time predicted their final behavioural accuracy. This latter result suggests that gaze dynamics reflect internal beliefs, and could help shed light on the computations that transform visual perception to action in naturalistic settings.

## RESULTS

Monkeys and humans performed a visual navigation task in which they used a joystick to steer to a cued target location in a three-dimensional virtual reality (VR) environment without allocentric reference cues (*i.e.* stable landmarks) (Fig 1A, **Methods**). At the beginning of each trial, a circular target blinked briefly at a random location within the field of view on the ground plane, and then disappeared. We gave subjects a joystick that controlled forward and angular velocities, allowing them to steer freely in two dimensions (Fig 1B). The subjects’ goal was to steer towards the target, and stop when they believed their position fell within a circular reward zone centered on the target. They received feedback about their performance immediately at the end of each trial.

**Figure 1.**
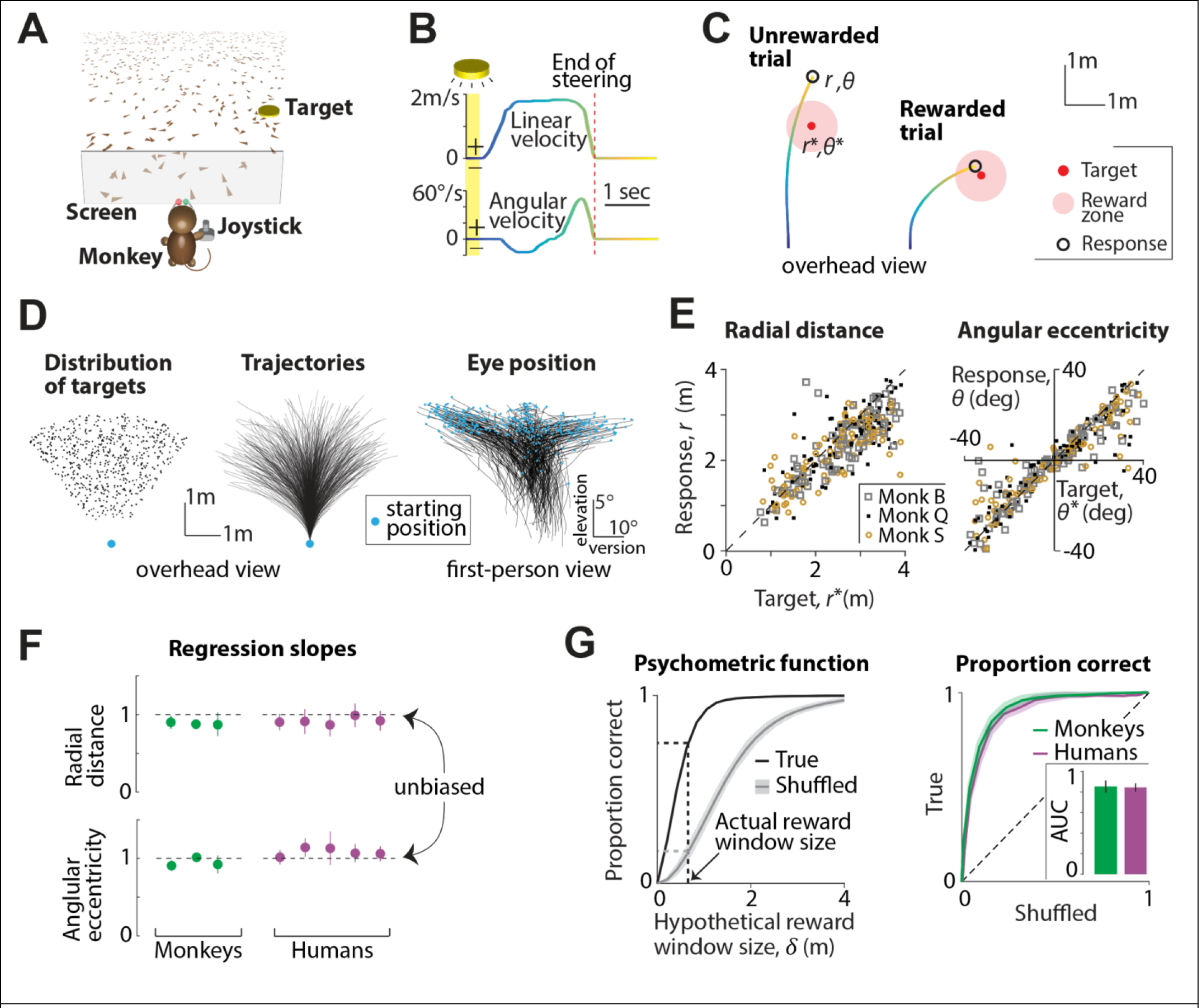
Primates can navigate by integrating optic flow. **A.** Monkeys and human subjects use a joystick to navigate to a cued target (*yellow disc*) using optic flow cues generated by ground plane elements (*brown triangles*). The ground plane elements appeared transiently at random orientations to ensure that they cannot serve as spatial or angular landmarks (**Methods**). **B.** The time-course of linear (*top*) and angular (*bottom*) velocities during one example trial. Yellow shaded region corresponds to the time period when the target was visible on the screen. Time is also coded by color. **C.** Example trials showing incorrect (*left*) and correct (*right*) responses of a monkey. Note that subjects had to stop within the reward window (0.6m for monkeys; adaptive window for humans, see **Methods**) to receive reward. **D.** *Left*: Overhead view of the spatial distribution of target positions across trials. Positions were uniformly distributed within subjects’ field of view. The actual range of target distances and angles was slightly larger for human subjects (**Methods**). *Middle*: Movement trajectories of one monkey during a representative subset of trials. Orange dot denotes starting location. *Right*: First-person view of the trajectories of eye movements (average of the two eyes) during the same trials. Abscissa and ordinate show horizontal version and elevation of the eyes respectively. Orange dots represent the initial eye position (when the target was turned OFF) on each trial. **E.** *Left*: Comparison of the radial distance of the monkey’s response (stopping location) against radial distance of the target across all trials. *Right*: Angular eccentricity of the response vs target angle. Black dashed lines have unity slope (unbiased performance). The subject’s starting location was taken as the origin. **F.** Subjects’ multiplicative biases in radial distance (*top*) and angular eccentricity (*bottom*) were quantified as the slopes of the corresponding linear regressions and plotted for individual monkeys (*green*) and human subjects (*purple*). Horizontal dashed lines denote the value of the slope that corresponds to unbiased behaviour. Error bars denote ±1 SEM across trials. **G.** *Left*: The proportion of correct trials of one monkey for various values of hypothetical reward window size (black). Shuffled estimates are shown in gray. *Right*: ROC curves for all subjects, obtained by plotting their true proportion of correct trials (from unshuffled data) against the corresponding chance-level proportions (from shuffled data) for a range of reward windows. Shaded area denotes standard deviation across subjects. Inset shows the average area under the curve (AUC) for monkeys (*green*) and human subjects (*purple*). See also **Figure S1-S2**.

Monkeys were first trained extensively using a staircase procedure (see **Methods**) until their performance stopped improving. Here, we will focus only on their post-training behaviour. At this point, the radius of the reward zone was fixed across trials (see **Methods**) and they received feedback in the form of juice reward at the end of the trial for correctly stopping within this zone (Fig 1C). In contrast, human subjects received no prior training on this task. Instead, we used an adaptive feedback scheme in which the radius of the reward zone was dynamically scaled using a staircase procedure to match individual subjects’ abilities (**Fig S1A**, see **Methods**). In practice, it took less than fifty trials for the performance of humans to stabilize (**Fig S1B**). Therefore, we ignored the first fifty trials collected from human subjects and focused our analyses on the remaining data.

Target locations were uniformly distributed at random over the ground plane area within the subject’s field of view (Fig 1D – *left*). The stimulus was nearly identical for both species except for minor details such as the range of target distances and the duration for which the target was visible (see **Methods**). All subjects were head-fixed, and we recorded each subject’s movement trajectory (Fig 1D – *middle*) as well as eye position (Fig 1D – *right*) throughout each trial.

### Behavioural performance

Figure 1E shows the performance of the monkeys in this task. Both radial distance (Fig 1E - *left*) and angular eccentricity (Fig 1E - *right*) of the monkeys’ responses (stopping location) were highly correlated with the target location across trials (*n* = 3 monkeys, Pearson’s *r* ± standard deviation, radial distance: 0.72 ± 0.1, angle: 0.84 ± 0.1) suggesting that their behaviour was appropriate for the task. To test whether their performance was accurate, we regressed their responses against target locations. The slope of the regression was close to unity both for radial distance (mean ± standard deviation = 0.92 ± 0.06) and angle (0.98 ± 0.1) suggesting that the monkeys were nearly unbiased (Fig 1F – *green*). We did notice modest undershooting for distant targets, an effect that is likely due to growing position uncertainty described in previous work (Lakshminarasimhan et al., 2018).

We showed previously that humans are systematically biased when performing this task without feedback, and that the bias was likely due to prior expectations that make them underestimate their movement velocities (Lakshminarasimhan et al., 2018). Consistent with those findings, human subjects overshot the target in an initial block of trials in which no feedback was provided (**Fig S1C**; *n* = 5, mean slope ± standard deviation, radial distance: 1.21 ± 0.2, angle: 1.78 ± 0.3), to a degree that was proportional to target distance. With feedback, however, the same subjects quickly adapted their responses to produce nearly unbiased performance (Fig 1F – *purple*, see **Fig S1D** for individual trials; mean slope ± standard deviation, radial distance: 0.95 ± 0.1, angle: 1.15 ± 0.2). Notably, this improvement in performance was maintained in a final block of trials in which feedback was withheld (**Fig S1E-F**; radial distance: 1.03 ± 0.15, angle: 1.2 ± 0.2) suggesting that learning of this task was stable. To maintain consistency with monkey data, we only consider human subjects’ data collected during the block of trials with feedback in the remainder of this work.

We wanted to know whether humans and monkeys had comparable accuracies. Because we used a slightly larger range of target distances for humans (see **Methods**), we could not directly compare the mean error magnitude of the subjects as it does not take differences in task difficulty into account. Instead, we used an approach that is conceptually similar to receiver operating characteristic (ROC) analysis to objectively compare the performance of monkeys and human subjects on a common scale. For each subject, we constructed a ‘psychometric function’ of performance as a function of hypothetical reward window size (Fig 1G; see **Methods**). By plotting the true psychometric function against one obtained by shuffling target locations across trials, we obtain the subject’s ROC curve. Chance-level performance would correspond to an area under the ROC curve (AUC) of 0.5, while perfectly accurate responses (zero error) will yield an AUC of one. The AUCs for both monkey and human subjects were quite large and statistically indistinguishable (mean ± standard deviation, monkeys: 0.85 ± 0.03, humans: 0.84 ± 0.05; *t*-test: *p* = 0.41) suggesting that they performed comparably. We emphasize that the individual visual elements comprising the ground plane were transient and could not be used as landmarks, so the performance of monkeys and human subjects in this task reflects their ability to integrate optic flow, rather than their ability to visually track a ground plane element. While it is still possible to avoid integrating optic flow by learning the precise sensorimotor transformation implemented by the joystick controller, we previously showed that the variability of human subjects is greatly affected by removing optic flow cues (Lakshminarasimhan et al., 2018)(**Fig S2A**). Likewise, monkeys rapidly adapt their actions in response to gain changes of the joystick controller (**Fig S2B**). This suggests that both monkeys and humans use optic flow to perform this task.

### Pattern of eye movements

To understand the role of eye movements, we recorded the position of the subjects’ eyes while they performed the task. Figure 2A shows the vertical and horizontal eye positions of one monkey during an example trial. On this trial, we noticed saccades (eye movements exceeding 200°/s) before the target was turned off (henceforth called *start* of the trial) and around the time when the monkey stopped moving (*end* of steering), but not in-between. This pattern of saccade timing was evident across trials, as seen in the trial-averaged density of saccades (Fig 2B). Across all datasets from monkeys, the average frequency of saccades during the trial was significantly smaller than that during the inter-trial interval (mean saccade rate ± standard deviation, during trials: 0.5 ± 0.3 Hz, between trials: 0.9 ± 0.5 Hz; paired *t*-test: *p* = 0.02). We noticed a similar tendency among human subjects although the comparison was not statistically significant (**Fig S3A**; during trials: 0.8 ± 0.5 Hz, between trials: 1.4 ± 1 Hz; *p* = 0.11). This suggests that subjects actively suppressed saccadic eye movements while steering. Moreover, the velocity of eye movements during steering was generally low, with magnitudes well below 20°/s both in monkeys (Fig 2C; mean ± std.: 16.2 ± 2.1 °/s) and in humans (**Fig S3B**; 11.4 ± 3.2 °/s).

**Figure 2.**
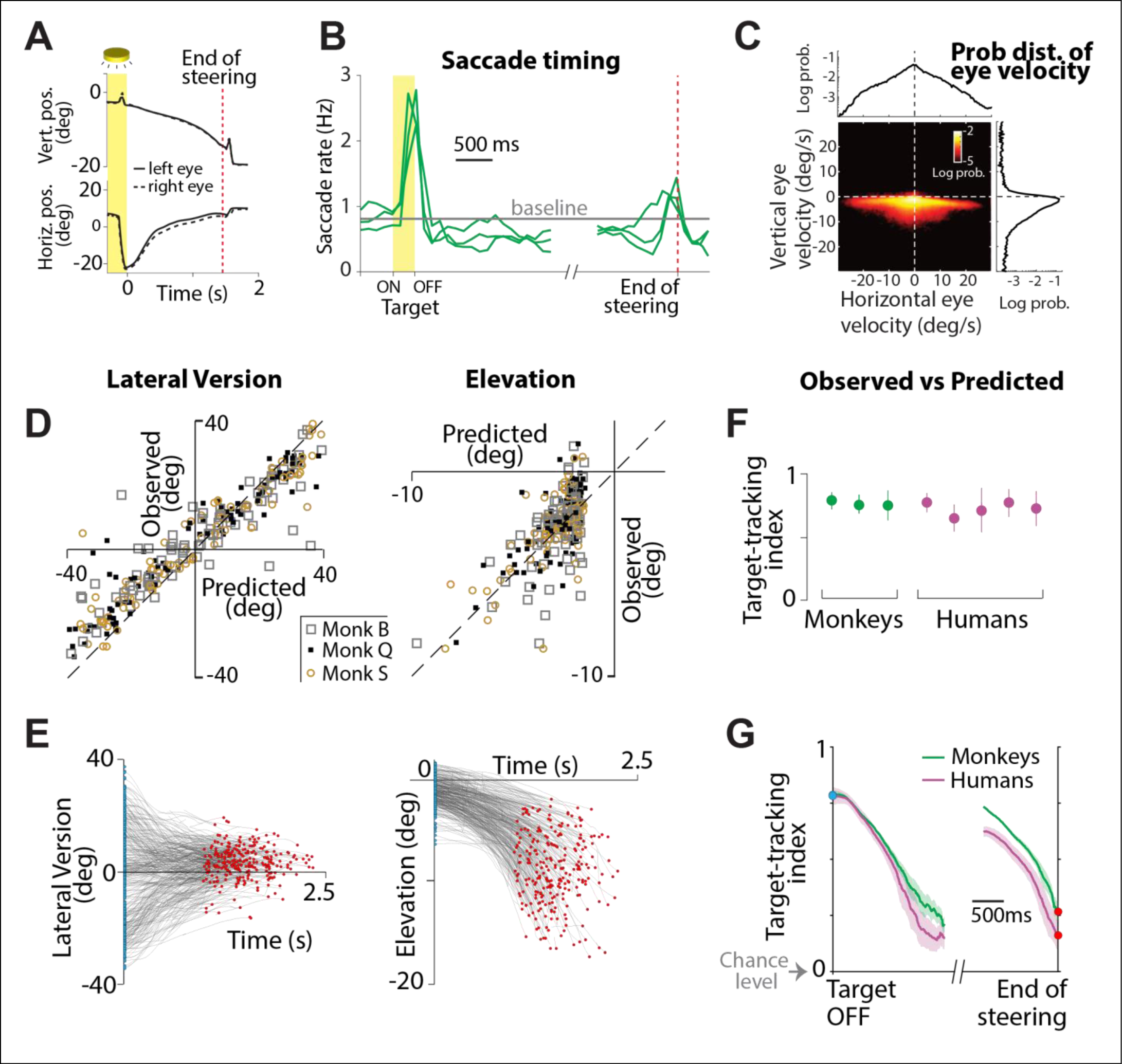
Eye movement dynamics during the task. **A.** Time-course of vertical (top) and horizontal (bottom) positions of the left (*solid*) and right (*dashed*) eyes of a monkey during one example trial (in degrees). Yellow region shows the period when a target was visible on the screen. Red dashed line corresponds to the end of steering in this trial. **B.** The time-course of the rate of saccades during the trial, averaged across all trials separately for each of the three monkeys. Trial-averaging was done by aligning trials relative to target onset (*yellow region*, before break on the x-axis) and end of steering (*red dashed line*, following the break). Grey line denotes mean saccade rate across monkeys during the period between trials. **C.** Joint probability density of the distribution over horizontal and vertical eye velocities, averaged across monkeys, while they steered towards the target. Marginals are shown in black. **D.** Comparison of the two major components (lateral version and elevation) of predicted and true eye positions in a subset of trials for all monkeys at the moment when the target was just turned OFF. **E.** Time-course of the two components (lateral version and elevation) of eye positions during a random subset of trials taken from all monkeys. Blue and red dots denote the times at which the target was turned OFF and the end of steering, respectively. **F.** Target-tracking index (defined in text) when the target turned OFF for individual monkeys (*green*) and humans (*purple*). Error bars denote ±1 SEM obtained either by averaging across recording sessions (for monkeys) or bootstrapping (for humans). **G.** Time-course of the target-tracking index, averaged across monkeys (*green*) and humans (*purple*). Grey arrow denotes the chance level tracking-index verified by shuffling procedure. Shaded region denotes ±1 SEM across datasets. See also **Figures S3**-**S8**.

Because saccades were mostly confined to periods when the animal was not actively steering and subjects appeared to make slowly-varying eye movements while steering, we reasoned that they may be continuously ‘tracking’ the (invisible) target with their eyes while they navigated to it. Note that as one steers towards the target location, the target would become progressively less eccentric and move downward in the visual field. Therefore, if subjects were to track the target, the magnitude of lateral version would tend towards zero and the eye elevation would become more negative with time (**Fig S4A**).

To quantitatively test whether subjects were tracking the target, we first generated ground truth theoretical predictions for the binocular position of their eyes during each trial, assuming that they maintained fixation at the center of the target throughout the trial (**Fig S4B**; **Methods** – Equation 1). We then compared this *prediction* against the *observed* eye position of the subject by expressing both quantities in terms of three standard components – lateral version, elevation and vergence (**Fig S4C**; see **Methods**).

We expect subjects’ eyes to be drawn to the target when it appears on the screen. So, at the very least, the theoretical predictions should be precise at trial onset. Indeed, the model predictions were highly correlated with the measured values of lateral version (Fig 2D – *left* and **Fig S5A** – *left*; Pearson’s *r* ± standard deviation, monkeys: 0.91 ± 0.1, humans: 0.85 ± 0.1) as well as elevation (Fig 2D – *right* and **Fig S5A** – *right*; monkeys: 0.60 ± 0.2, humans: 0.42 ± 0.2) at the beginning of the trial. The somewhat lower correlations for the latter are understandable because it is difficult to precisely fixate at the elevations for distant targets since they subtend a smaller visual angle. We verified this effect using simulations (**Fig S6**). Next, we examined the time-course of eye movements during the trial and found a striking qualitative correspondence to the predicted dynamics (Fig 2E, **S5B**): as the trial progressed, lateral version became increasingly more concentrated around zero (**Fig S5C –** *left*) while eye elevation was significantly lower (**Fig S5C –** *right*). The correlation between predicted and observed values remained significantly greater than zero throughout the trial for both components (**Fig S5D**). This is quite remarkable because the target appeared only transiently at the beginning of the trial.

On the other hand, the correspondence between predicted and observed vergence was less clear. Doing this comparison for our task was challenging because about 90% of the full range of vergence angles is known to occur within gaze distances below one meter (Howard, 2012) and the predicted change in vergence is negligible for gaze distances beyond 2m (**Fig S4C –** *bottom right*). Only two of the three monkeys exhibited vergence values that weakly correlated with the predictions at trial onset (**Fig S7A**) and a tendency to make convergent eye movements as they approached the target (**Fig S7B**), an effect that was also absent in human subjects (**Fig S7B-D**). It is possible that this inconsistency is due to the previously documented difficulty in executing voluntary vergence movements to imagined moving targets (Erkelens et al., 1989). Moreover, this difficulty is likely exacerbated in VR where vergence eye movements must be executed without changing accommodation to maintain a clear retinal image of onscreen objects (Hoffman et al., 2008; Lambooij et al., 2009; Shibata et al., 2011). Therefore, we did not consider the vergence component for further analyses.

To quantify the extent to which a subject’s eyes tracked the target, we expressed the eye position as a two-dimensional vector comprised of lateral version and elevation, and computed a *target-tracking index* that measures how precisely the subjects’ eyes tracked the target. Specifically, this quantity was given by the square root of the fraction of variance in the observed eye position that was explained by the prediction (**Methods** – Equation 2). An index of one implies that the subject consistently looked at the center of the target, while zero denotes lack of correspondence between target and gaze locations. The target-tracking index was quite high at trial onset (during the first 500ms) when the target had just disappeared (Fig 2F; mean ± standard deviation, monkeys: 0.73 ± 0.05, humans: 0.71±0.05). Although this slowly dropped during the trial, the index at the end of the trial (during the last 500ms) remained well above zero (Fig 2G; mean ± standard deviation, monkeys: 0.35 ± 0.1, humans: 0.18±0.05), implying that subjects tend to maintain gaze at the target location while they steer towards it. Alternative measures comparing observed eye positions to the predictions exhibited qualitatively similar dynamics, so the above result is robust to the precise definition of the target-tracking index used here. While the above analysis quantifies the extent of correlation between gaze and target location, it does not specify the timescale of those correlations. To estimate this timescale, we analyzed the cross-correlogram between them and found that subjects’ eye positions did not systematically lead or lag the predictions based on the contemporaneous target location (**Fig S8**). This suggests that eye movements were not predicting future target locations, although the computations used to estimate the target location could still be predictive.

The tracking index quantifies how subjects’ dynamical state is encoded in their continuous-valued eye position. However, recent work has highlighted the importance of discrete saccadic eye movements in mediating flow-tracking behaviour of humans and primates (Knöll et al., 2018). Therefore, we wanted to know whether saccades aided the target-tracking behaviour of our subjects. To test this, we first compared the distribution of saccade amplitudes during three non-overlapping epochs of the experiment – inter-trial periods when saccades tend to be exploratory, target-presentation phase when saccades are guided by the external stimulus, and the ensuing task phase when subjects steered using optic flow (Fig 3A). We found that across monkeys, the amplitude of saccades was much lower during the task phase than during other epochs (Fig 3B; mean ± SE: inter-trial – 10.2 ± 1.6°, target-presentation – 14.4 ± 2.2°, task phase – 7.1 ± 1.2°) suggesting that saccades made while steering may serve to correct errors in tracking the target. To directly test this, we computed a saccade-triggered average of the target-tracking error and found that this error dropped significantly (peak decrease of 2.4 ± 0.4°; *p* < 10^−10^, *t*-test) shortly after saccade onset (Fig 3C). Following Knoll et al. (2018), we used regression analysis to determine the precise relationship between saccade amplitude and the dynamics of target-tracking error (**Methods**). The amplitude of both vertical and horizontal components of the saccade were influenced by tracking error during the previous 200ms suggesting that these saccades were indeed made in the direction of the target (Fig 3D). Moreover, the regression kernels were biphasic implying that the saccades overcompensated for the tracking errors. Finally, if these saccades were corrective, they should rely on the subjects’ internal estimate of the target location making them increasingly unreliable over time due to the buildup of uncertainty. Indeed, the strength of the regression kernel was weaker for later saccades (Fig 3E; peak-to-peak difference in weights for vertical component: first saccade – 0.44 ± 0.1, third – 0.20 ± 0.15; horizontal component: first saccade – 1.5 ± 0.4, third – 0.15 ± 0.2), thereby signaling a drop in saccadic precision over time. This suggests that these saccades were not stereotyped reflexive responses but were dynamically modulated by ongoing cognitive computations analogous to “catch-up” saccades typically observed during smooth pursuit of visible targets (Daye et al., 2014a; Orban de Xivry et al., 2008). Although the amplitude of saccades made by human subjects were not significantly smaller while steering (**Fig S10A**), regression analysis revealed a strong association between tracking error and saccade amplitude but with slightly shorter integration windows. As observed for monkeys, the strength of this association was lower for saccades that happened later (**Fig S10B**) reflecting a potential influence of noisy integration.

**Figure 3.**
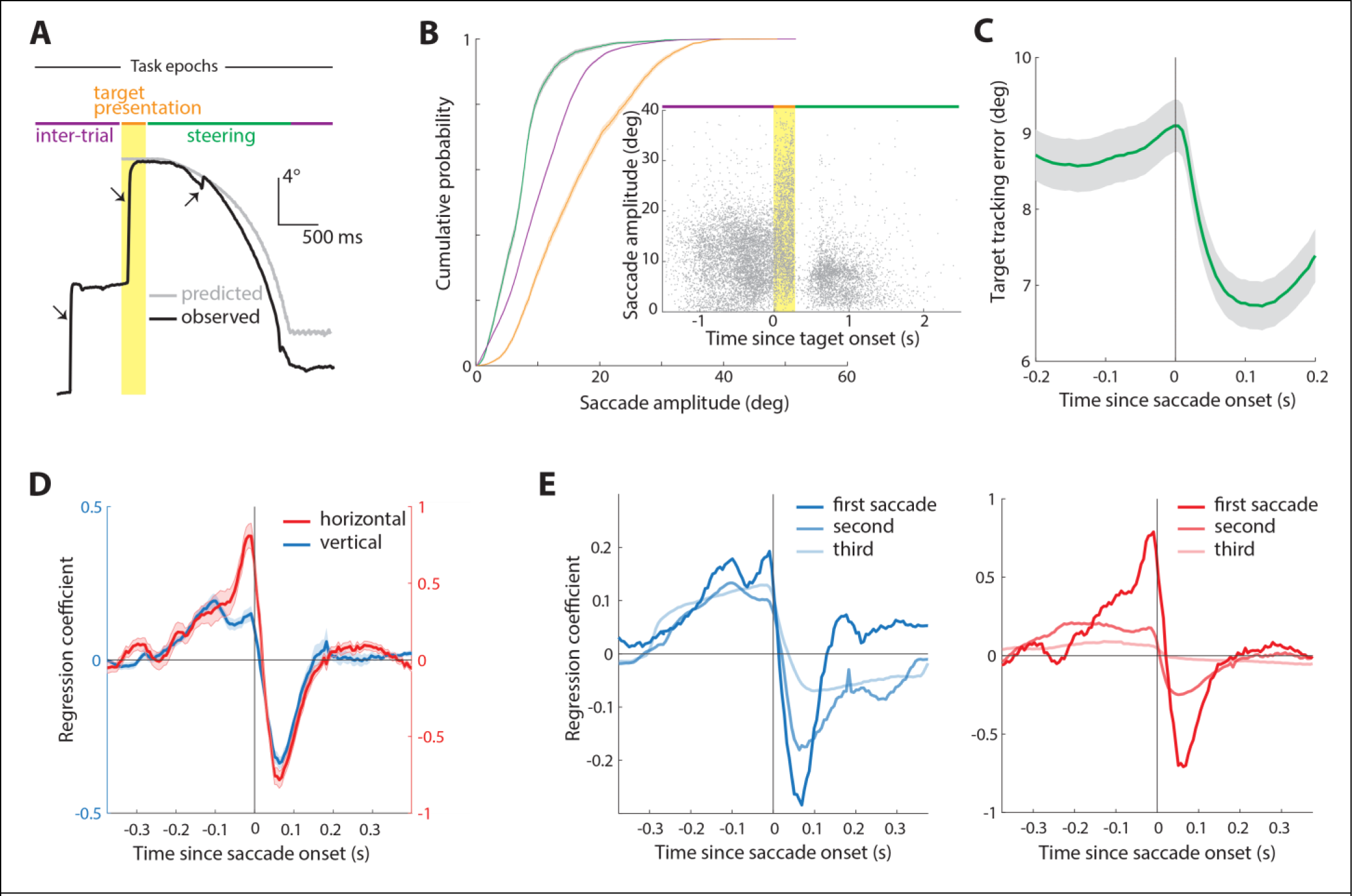
Saccadic eye movements aid target tracking. **A.** Time-course of observed (black) and predicted (gray) vertical position of the eyes of a monkey. Black arrows indicate saccades made during three different task epochs (inter-trial, target presentation, and steering periods). Yellow region shows the period when a target was visible on the screen. **B.** Empirical cumulative distribution function of saccade amplitudes conditioned on the task epoch, averaged across monkeys. Inset shows amplitudes of individual saccades as a function of their timing. **C.** Average saccade-triggered target tracking error during a time-window around saccades made during steering. **D.** The time-course of coefficients obtained by linearly regressing the amplitudes of the two components of saccades (blue – vertical, red – horizontal) against the corresponding components of the target tracking error (**Methods**). **E.** Similar to D, but showing coefficients for regression done separately for the first, second, and third saccades made during steering (left – vertical component, right – horizontal component). See also **Figure S10**.

### Eyes convey internal beliefs about target

Subjects could not have possibly been tracking the observed target location, since the target disappeared at the beginning of the trial. A plausible explanation for their pattern of eye movements is that subjects tracked the location at which they *believed* the target was present. As they integrate their movements, subjects need to continuously update their internal estimate of the relative goal location, and perhaps their eye movements reveal those estimates. If this is the case, then we should be able to better predict their eye position when their beliefs are more accurate. We tested this both across subjects and across trials within each subject.

To test this across subjects, we used the variability in subjects’ stopping positions to first quantify the level of uncertainty in their position estimates (**Methods**). Due to the low trial count of individual human subjects, we pooled trials from all humans into a single dataset. Because uncertainty in knowing one’s location should limit one’s ability to visually track the target, we used the estimated uncertainties to calculate an approximate *upper bound* on the target-tracking index for each dataset (Fig 4A, **Methods** – equations 3). This upper bound serves to capture the heterogeneity in the spatial profile of uncertainty both across subjects (Fig 4B – *left*) and across sessions within each monkey (**Fig S11A**). Across all datasets, the target-tracking index observed towards the end of the trial (during the last 500ms) was weakly but significantly correlated with the theoretical upper bounds (Fig 4B – *right*; Pearson’s *r* = 0.26, *p* = 0.029). This suggests that differences in the ability to track the target with the eyes is due, at least in part, to differences in the magnitudes of positional uncertainty between subjects.

**Figure 4.**
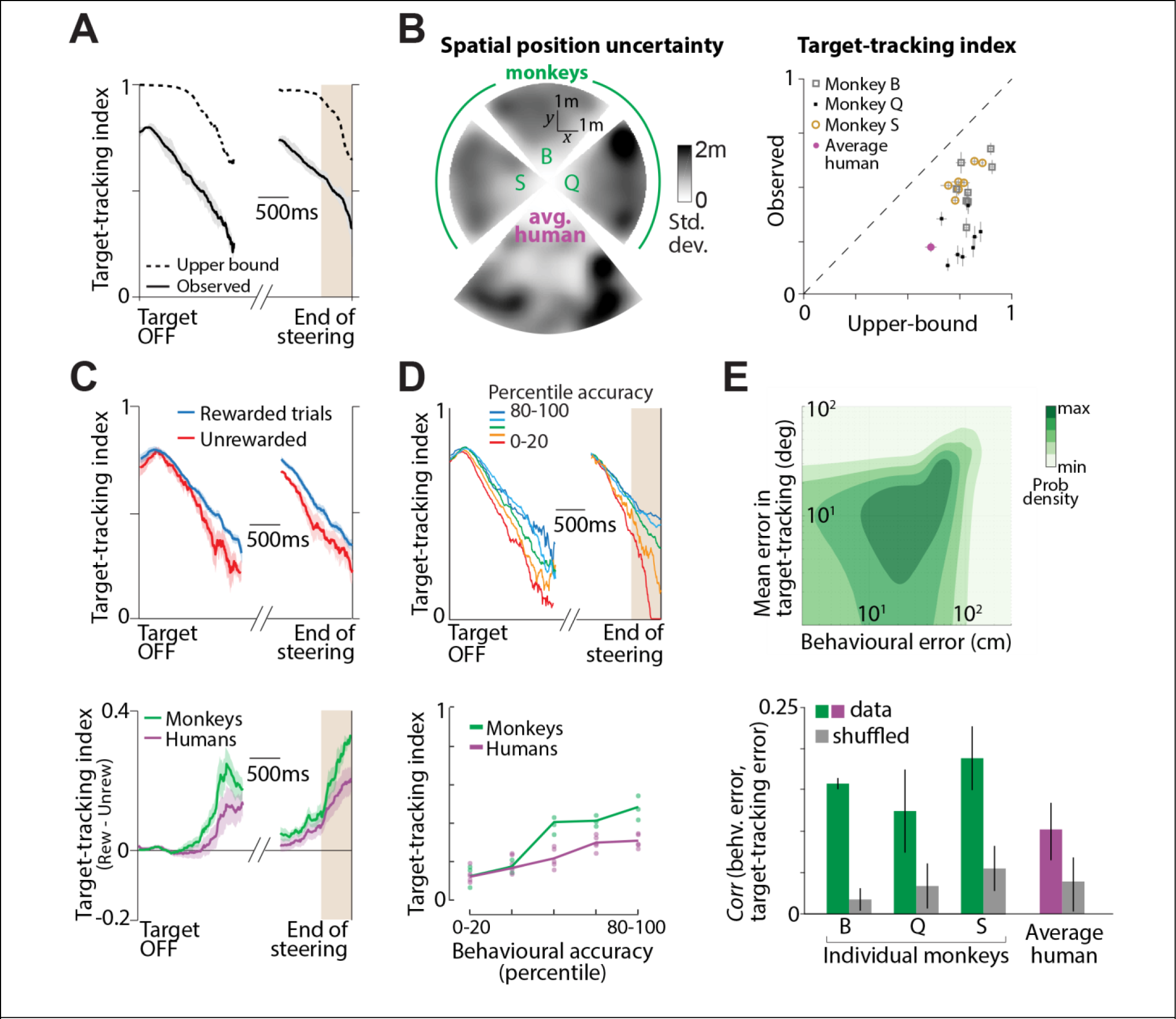
Accurate target-tracking is associated with increased task performance. **A.** Time-course of the target-tracking index for an example session computed using a monkey’s actual eye movements (*black solid*) and its theoretical upper-bound (*black dashed*) determined using variability in the monkey’s behavioural response (**Methods**, equations 3**-4**). **B.** *Left*: Overhead view of the spatial map showing the standard deviation of stopping positions as a function of target location for individual monkeys and the average human subject. The maps of monkey S & Q, and of the humans, have been rotated solely for visualization. All subjects shared the same range of target angles (±40°) and distances (up to 4m for monkeys, 6m for humans). *Right*: Comparison of the observed target-tracking index against the corresponding theoretical upper bound (averaged over the last 500ms of the trials) across all individual datasets. Trials from human subjects were pooled together (see text for explanation). Dashed line has unity slope and error bars denote ±1 SEM obtained by bootstrapping. **C.** *Top*: Time-course of the target-tracking index for one example monkey shown separately for trials in which he stopped within the reward zone (0.6m from the target, *blue*), or stopped outside it (*red*). Shaded regions denote ± 1 standard error estimated by bootstrapping. *Bottom*: The difference between tracking coefficients during rewarded and unrewarded trials for all subjects (monkeys in green, humans in purple). For human subjects, trials in which the subject’s final position was within 0.6m of the center of the target were considered ‘rewarded’ for the purpose of classification. **D.** *Top*: We divided trials into five groups depending on the magnitude of the subject’s error *i.e.*, final (stopping) distance to the target. Time-courses of the target-tracking index are shown for the five trial groups for one monkey (*dark blue*: most accurate; *dark red*: least accurate). *Bottom*: Average value of the target-tracking index during the final 500ms before end of steering (brown shaded region in the top panel) as a function of percentile accuracy for all individual subjects. Solid lines show average across subjects. Across all subjects (humans and monkeys), there was a significant correlation between accuracy and tracking coefficient (Pearson’s *r* = 0.68, *p* = 3.1 × 10^−5^). **E.** *Top*: Joint distribution of the behavioural error and the target-tracking error across trials of one recording session from one monkey. *Bottom*: The mean correlation between behavioural and target-tracking errors of individual monkeys before (green) and after (gray) a shuffling procedure to control for the effects of trial difficulty (see text). Error bar denotes ±1 SEM obtained by bootstrapping. See also **Figure S11**.

We also tested whether eye movements reflect fluctuations in the subject’s belief about their location across trials. Because subjects were more precise during rewarded (**Fig S11B –** *left*) than during unrewarded trials (**Fig S11B –** *middle*), we expect subjects to track the target more accurately during rewarded trials (**Fig S9B** – *right*). We computed the target-tracking index separately for the two groups of trials and found that it was indeed higher during rewarded trials (Fig 4C – *top*). The difference between the target-tracking indices during the two sets of trials grew as the trial progressed, and was significantly greater than zero at the end of the trial (Fig 4C – *bottom*; mean difference ± standard deviation during the period shaded in grey – monkeys: 0.19 ± 0.05, *p* = 4.8 × 10^−3^; humans: 0.13 ± 0.05, *p* = 3.1 × 10^−2^; bootstrap test, 10,000 bootstrap samples). In fact, when trials were stratified based on behavioural accuracy, we found that the tracking index increased with behavioural accuracy (Fig 4D). To more directly test whether there was a fine-grained relationship between eye movements and task performance, we estimated the correlation between the behavioural error (distance between the stopping location and the target) and the target-tracking error (mean absolute difference between the actual eye position and the theoretical prediction, see **Methods**) across trials (Fig 4E – *top*). To control for possible spurious effects of trial difficulty, we computed a shuffled estimate by subdividing the trials into groups based on initial target distance and then shuffling the trials within each group (see **Methods**). We found that the behavioural and target-tracking errors were significantly correlated across trials (Fig 4E – *bottom*, Pearson’s *r* ± standard deviation across all datasets – true: 0.14 ± 0.04, controlled shuffle: 0.04 ± 0.02; *p* = 9.1 × 10^−3^, paired *t*-test) further reinforcing the view that subjects are tracking their internally estimated goal location with their eyes.

### Purely reflexive eye movements do not explain target-tracking behaviour

In principle, the above results could also be produced by purely reflexive eye movements, driven solely by optic flow (ocular following response or OFR). For instance, if subjects’ eye velocity is perfectly correlated with their perceived movement velocity, then oculomotor errors would be proportional to perceptual errors, potentially explaining the relatively poor target-tracking in erroneous trials. However, past studies have shown that errors in reflexive eye movements are uncorrelated with perceptual errors (Blum and Price, 2014; Boström and Warzecha, 2010; Glasser and Tadin, 2014; Price and Blum, 2014) suggesting that the observed eye movements may not be entirely reflexive. Two further pieces of evidence in our own monkey data support this.

First, in a subset of sessions we recorded the stimulus movie of the complete block of trials and replayed them back to the animal at the end of the session, but with the joystick withheld (see **Methods**). All aspects of the task structure during this replay block were identical to the initial block of trials (e.g. the monkey still received juice reward at the end of the corresponding trials), except the animal only viewed a movie of the stimulus rather than actively performing the task. Importantly, monkeys were still free to move their eyes. In general, eye movements were weaker during passive viewing than during active task (**Fig S12A, B**). Across monkeys, the magnitude of eye velocity was much smaller during passive block even though both blocks had identical visual stimuli (**Fig S12C**). We analysed the target tracking behaviour by computing the target-tracking index separately for the two blocks of trials. Figure 5A (top panel) shows the time-course of the target-tracking index of one monkey during the both blocks of trials. In this monkey, the tracking index was much lower during passive viewing (red vs blue). Because OFR is, by definition, involuntary and difficult to suppress, this suggests that eye movements contributing to the high target-tracking index during active steering must have been voluntary. Note however that the tracking index during passive viewing is poor right from trial onset, perhaps because the monkey did not consistently look at the target initially when it appeared on the screen. We wanted to know whether OFR dynamics, coupled with the appropriate boundary condition (looking at the target when it initially appears) might be sufficient to give the impression that the animal is tracking the target. We simulated this model by shifting the initial eye position on each trial of the passive block to match the corresponding trial in the active block, a procedure that left the eye movement dynamics unaltered (Fig 5A – *black*). The tracking index of this simulated model was substantially lower than that observed during the active block of trials, suggesting that the target-tracking behaviour is likely to be a voluntary response. In all three monkeys, the target tracking during the active task was significantly stronger than during either the passive viewing condition or the OFR model (Fig 5A – *bottom*; mean difference ± standard deviation during the period shaded in brown, active: 0.27 ± 0.1, passive: 0.08 ± 0.1, OFR model: 0.07 ± 0.1; *p* < 0.01, bootstrap test). The difference between conditions was small in one monkey (labelled ‘Q’ in Fig 5A – bottom; **Fig S12** – rightmost), possibly because this animal was mentally performing the task even during passive viewing.

**Figure 5.**
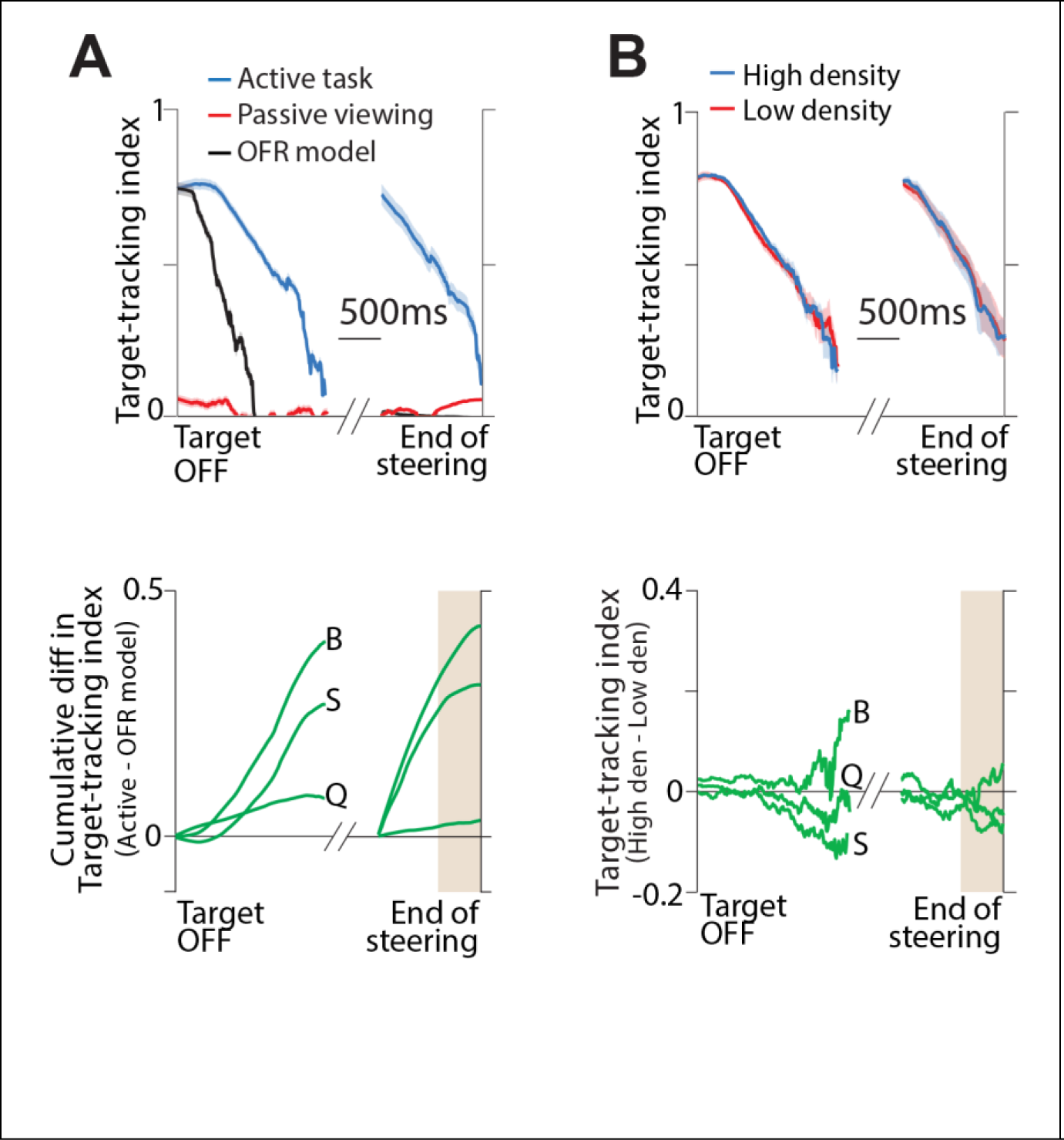
Steering-induced eye movements are not reflexive. **A.** *Top*: Time-course of target-tracking index for one monkey during trials in which he performed the task (*blue*) or passively viewed the stimulus identical to the one generated when performing the task (*red*). Black trace shows the tracking index of an OFR model simulated by translating the initial eye position on each passive trial to match the corresponding active trial (see text). Tracking indices at time points with negative variance explained were clipped to zero. *Bottom*: The time-course of the cumulative difference between the target-tracking index on active trials and the simulated OFR model for individual monkeys. **B.** *Top*: Time-course of the tracking index of one monkey during trials in which the density of ground plane elements was either high (*blue*) or low (*red*). *Bottom*: The difference between target-tracking index under high and low density conditions for individual monkeys. Brown shaded regions in the bottom panels correspond to the 500ms time-window considered for statistical testing. See also **Figure S12**.

Second, OFR is known to be sensitive to signal strength (Barthelemy et al., 2009; Quaia et al., 2012). To test whether target tracking depends on signal strength, we manipulated stimulus reliability by randomly interleaving trials with two different densities of ground plane elements by more than an order of magnitude (see **Methods**). We analysed the two sets of trials separately, but found no significant difference between the target-tracking index (Fig 5B; mean ± standard deviation across subjects, low density: 0.28 ± 0.1, high density: 0.31 ± 0.1). Therefore, the pattern of eye movements observed during this task likely represent volitional movements, rather than reflexive ones.

### Inhibiting eye movements worsens task performance

Since eye movements were predictive of subjects’ navigational performance, we wanted to know if they were essential for performing the task. To test this, we asked five human subjects to perform a variation of the task in which we overlaid a cross on top of the target location and instructed them to fixate on this cross for as long as it appeared on the screen. In half the trials (‘Eyes-moving’ condition), the fixation cross disappeared along with the target so that subjects were free to produce eye movements as before. In the remaining trials (‘Eyes-fixed’ condition), the cross remained at the same location on the screen throughout the trial and subjects had to perform the task without moving their eyes (see **Methods**). Although we did not penalize subjects for breaking fixation, we verified offline that they maintained fixation as instructed (Fig 6A and **Fig S13**). We assessed their behavioural performance by comparing the area under the ROC curve (AUC), and found that performance was significantly impaired in the ‘Eyes-fixed’ condition (Fig 6B; *n* = 5 humans, mean AUC ± standard deviation; Eyes-moving: 0.85 ± 0.07, Eyes-fixed: 0.77 ± 0.07, *p* = 2.5 × 10^−3^, paired *t*-test). Figure 6C shows the responses of individual subjects.

**Figure 6.**
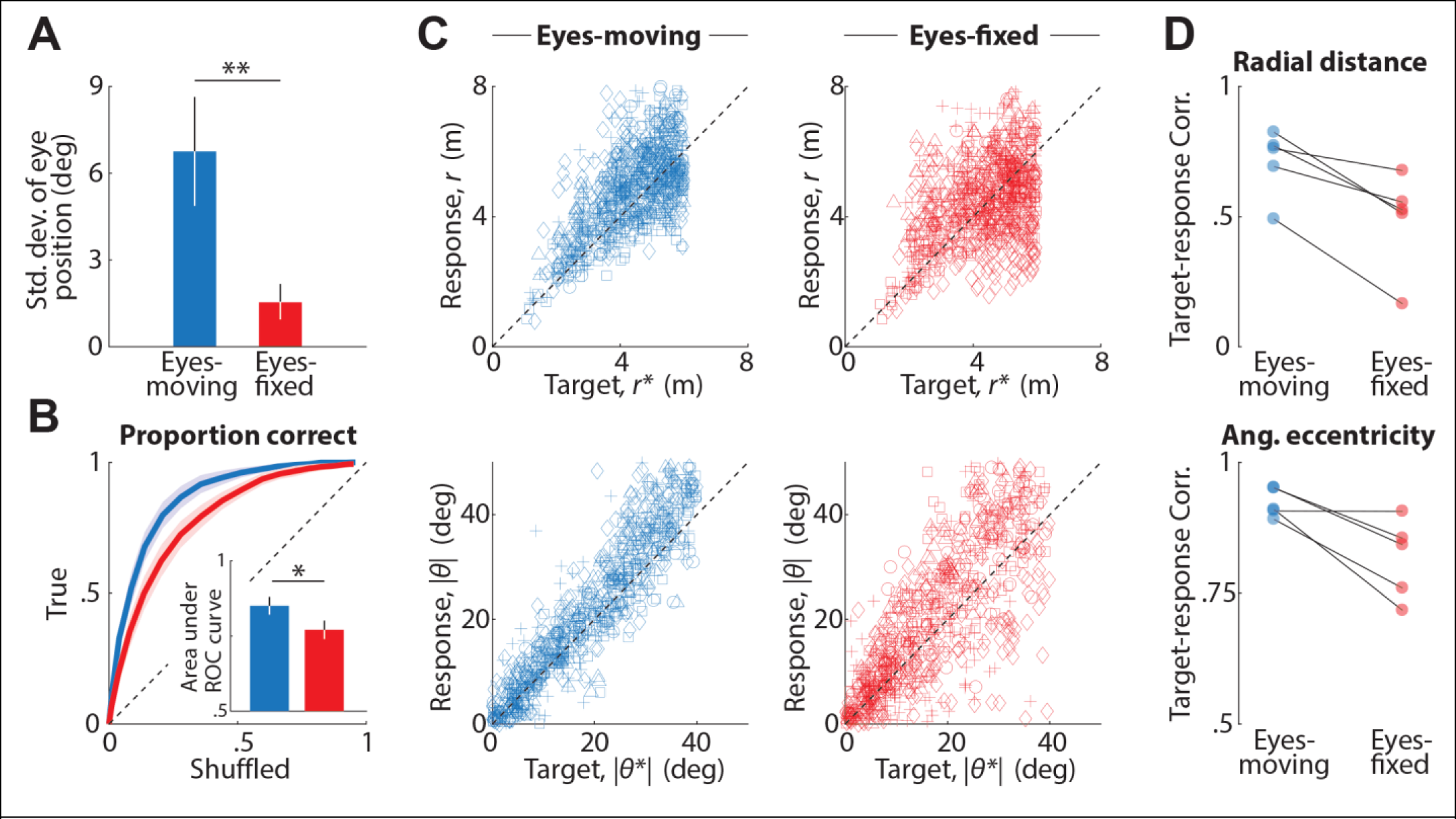
Fixation affects task performance. **A.** Trial-averaged temporal variability of subjects’ eye position, quantified by standard deviation (see **Methods**) during ‘Eyes-moving’ (*blue*) and ‘Eyes-fixed’ (*red*) trials. Error bars denote standard deviation across subjects (** *p* = 1.2 × 10^−3^, paired *t*-test). **B.** ROC curves averaged across subjects, for trials in the ‘Eyes-moving’ (*blue*) and the ‘Eyes-fixed’ condition (*red*). Inset shows the area under the two curves. Error bars denote standard deviation across subjects (* *p* = 2.5 × 10^−3^, paired *t*-test). **C.** *Top*: Comparison of the radial distances of the response and the target on trials in ‘Eyes-moving’ (*blue*) ‘Eyes-fixed’ (*red*) conditions. Different symbols denote different human subjects. *Bottom*: Comparison of the (absolute) angular eccentricity of the response and target under the same two conditions. **D.** *Top*: Pearson’s correlation coefficient between the radial distance of subjects’ response and the target under ‘Eyes-moving’ (*blue*) and ‘Eyes-fixed’ (*red*) for all individual subjects. *Bottom*: Similar comparison for the absolute angular eccentricity of target and response under the two conditions. See also **Figure S13**.

Although subjects were nearly unbiased under both conditions, the correlation between target and response locations was significantly lower in the absence of eye movements (Fig 6D; mean ± standard deviation; *Corr*(*r*, *r*^*^), Eyes-moving: 0.71 ± 0.1, Eyes-fixed: 0.49 ± 0.2, *p* = 0.011, paired *t*-test; *Corr*(|*θ*|, |*θ*^*^|), Eyes-moving: 0.92±0.03, Eyes-fixed: 0.82±0.1, *p* = 0.035). These results suggest that subjects benefit when their eyes can track the internally estimated goal location in this task.

## DISCUSSION

Using a virtual visuomotor navigation task that requires continuous integration of self-motion, we showed that humans and monkeys execute similar eye movements while steering. By comparing the eye movement dynamics against predictions for a hypothetical agent that maintained fixation at the target, we demonstrated that subjects likely tracked the imagined target location when steering towards it. Although subjects tended to suppress saccades while steering, saccades were, on average, aimed towards the target and thus helped correct errors in target tracking. While the tracking index remained significantly above chance throughout the trial, it nonetheless decreased over time. This is expected because the target disappears, so subjects cannot directly measure its true position but must instead rely on an internal estimate computed by integrating optic flow with knowledge of the controller dynamics. We have previously shown that human subjects perform near-perfect integration in this task (Lakshminarasimhan et al., 2018). Nevertheless, due to noise in the integration process, the error in the internal estimate of target location on any given trial should grow over time. Consequently, even if those estimates are unbiased, their precision worsens, leading to a decrease in the target-tracking index (Fig 4A – dashed line). Consistent with this, the precision of error-correcting saccades gradually deteriorated as the trial progressed (Fig 3E). Therefore, the observed decrease in target tracking is an inevitable consequence of noisy observations and noisy integration, and in fact serves to expose the growing uncertainty in subjective beliefs.

### Inferring belief dynamics

The task design used in this study was motivated by the need to ultimately understand neural computations governing belief dynamics that transform sensory inputs to motor output. In the real world, these belief dynamics correspond to *subjective* estimates of latent-variable dynamics, thus making them difficult to measure and relate to neural activity. Classic experimental paradigms used with primates attempt to achieve this by training animals to provide discrete responses to simple stimuli, which reduces the dimensionality of state and action spaces, limiting their potential to shed light on natural computations. On the other hand, paradigms that let rodents loose in open arenas mimic natural behaviour at the expense of sacrificing control over the nature of computations they perform. One exception is a recent paper (Knöll et al., 2018) in which the authors used stimuli with rich spatiotemporal dynamics to elicit continuous oculomotor behaviour from primates with minimal reinforcement. While the Knoll et al. (2018) task is similar to ours in that it leverages unconstrained eye movements to remedy several shortcomings of historical approaches, the computation performed by the animals in their task is relatively straightforward: tracking a visible focus of expansion in optic flow. In contrast, we asked animals to integrate momentary information about their movements to navigate to a goal location without providing explicit position cues. This forces the animal to track the dynamics of a *latent* variable (position of the target relative to them) – a more challenging computation. Moreover, we trained our animals only by rewarding them for reaching the goal location. Eye movements were not explicitly reinforced, yet our post-hoc analysis revealed that the continuous-valued, time-varying eye position encoded subjective beliefs about the time-varying latent variable. We hope that this approach of covertly measuring belief dynamics will serve as a useful template for future studies.

### The nature of eye movements

While steering towards the target, subjects executed slow eye movements, and tended to suppress saccades. To understand the nature of slow eye movements, we analyzed three separate components of eye position: lateral version, elevation, and vergence. In all subjects, the dynamics of the first two components were smooth and consistent with the predicted dynamics for pursuing the invisible target. In contrast, only two of the three monkeys made convergent eye movements as they approached the target location. Vergence eye movements also did not show clear dependence on the target location in human subjects. Under natural conditions, vergence eye movements are typically evoked either by binocular disparity or by a need to accommodate to blurred visual stimuli (Horwood and Riddell, 2008; Howard, 2012). Accordingly, vergence responses to imagined targets are unreliable (Erkelens et al., 1989). Moreover, accommodation demands are somewhat unnatural in VR because objects on the screen all share the same focal length. In light of these limitations, it is not very surprising that we were unable to measure vergence eye movements that varied systematically with target position in all subjects.

By analyzing eye movements during stimulus playback, we ruled out the possibility that the smooth dynamics correspond to pure ocular following reflex (OFR) induced by optic flow. Because these eye movements were always preceded by fixating a visible target and occurred in parallel with computations for mentally tracking that same target, they are functionally more similar to smooth-pursuit eye movements. Despite ample evidence for smooth-pursuit eye movements in the absence of foveal stimulation in humans (Becker and Fuchs, 1985; Missal and Heinen, 2017; Wyatt et al., 1994) and rhesus macaques (Ilg and Thier, 1999), smoothly tracking a purely imaginary object is thought to be difficult (Spering and Montagnini, 2011). This is because, in the absence of dynamic information about target motion, the pursuit velocity gradually decays to zero (Barnes, 2008; Missal and Heinen, 2017). However, when the underlying model for target motion is known, subjects can use their dynamic internal representation of the target to make predictive smooth pursuit during target blanking (Adams et al., 2012; Orban de Xivry et al., 2013, 2008). In our task, the dynamics of optic flow completely determine the (relative) motion of the target and can subsequently drive eye movements. Furthermore, the flow fields were self-generated rather than simulated, a condition that has previously been shown to improve pursuit of occluded targets (Danion et al., 2017; Gauthier et al., 1988; Vercher and Gauthier, 1992). Finally, we note that a moderate contribution of OFR induced by optic flow cannot be completely excluded, so it is possible that the pattern of eye movements reported here is ultimately composed of a mixture of reflexive signals that encode velocity of self-motion and predictive signals that encode the internal estimate of relative target location.

Finally, saccadic eye movements, although infrequent, contributed to tracking the target. Moreover, the amplitude of these saccade was largely influenced by target-tracking error during the previous ~200ms. These results suggest that the mechanism responsible for generating saccades in this paradigm may be similar to the ones at play in flow-tracking (Knöll et al., 2018) and smooth pursuit of visible objects (Daye et al., 2014b; Orban de Xivry et al., 2008). One reason for the relatively low frequency of saccades in this study could be that motion in our task was self-generated and predominantly smooth, whereas saccades in smooth-pursuit experiments are primarily due to unexpected jumps in target velocity (de Brouwer et al., 2002).

### Possible function of tracking eye movements

The experimental task was specifically designed to ensure that subjects would attempt to *mentally* track the goal location by integrating momentary sensory evidence about movement provided by optic flow. In principle, this can be accomplished without *physically* tracking the believed goal location with one’s eyes. Yet we noticed a significant decline in task performance when eye movements were suppressed. This is consistent with previous results that demonstrated that real-world driving performance is impaired when eye movements are constrained (Wilson et al., 2008). Although this does not demonstrate a need to make tracking eye movements, it suggests that eye movements play an important role in neural computations for navigation. Indirect evidence of a role for slow eye movements in visually-guided navigation comes from a recent study of path integration, in which subjects used a joystick to reproduce previously-experienced self-motion (Churan et al., 2018). Eye movements during the reproduction phase were similar to those during initial exposure even when optic flow was removed. This suggested that eye movements constitute a form of mental imagery that, if suppressed, hampered memory retrieval (Johansson and Johansson, 2014; Johansson et al., 2012). Our findings extend this to naturalistic settings and argue that eye movements have a more dynamic role in path integration. The precise computational advantage of the specific eye movement dynamics observed in our task is unclear. Below, we propose two potential theories.

One possibility is that eye movements directed towards the intended goal location stabilizes the mental image of the goal, and could reduce the computational complexity of estimating self-motion from optic flow similar to the effect of foveal image stabilization (Lappe et al., 1999; Longuet-Higgins and Prazdny, 1980; Perrone and Stone, 1994; Sandini and Tistarelli, 1990; Sandini et al., 1986). Normative mathematical theories posit that maintaining gaze at a point on the intended path can greatly simplify the problem of exploiting optic flow (Glennerster et al., 2001; Kim and Turvey, 1999; Wann and Swapp, 2000). Therefore, the eye movements reported here may constitute a closed-loop visuomotor process in which subjects integrate sense data (optic flow) to dynamically update their beliefs about the relative goal location, and in turn, use them to guide future eye movements in order to acquire new sense data in a computationally useful format. In this view, eye movements primarily aid optic flow processing.

Alternatively, the observed eye movements might simply be an embodiment of subjects’ dynamically evolving internal beliefs about the goal. Humans have a well-documented tendency for externalizing their internal representations (Barsalou, 2008; Spivey, 2007), with eye movements sometimes employed as a pointing device to visible as well as invisible objects, much like one’s index finger (Ballard et al., 1995, 1997; Spivey and Geng, 2001). By allowing dynamic beliefs about the relative target location to continuously modulate eye movements in this task, the brain could piggyback on the oculomotor circuit and reduce the computational burden on working memory. Consistent with this interpretation, there is overwhelming evidence for decision-related responses in primate oculomotor brain areas (de Lafuente et al., 2015; Shadlen and Newsome, 1996), and such responses are thought to drive eye movements (Joo et al., 2016). Therefore, in this view, primates use gaze as an affordance to efficiently update and store the output of integrating optic flow.

Although the above accounts are not mutually exclusive, simultaneously recording the neural activity from the primate sensory, oculomotor, and decision areas during this task might shed light on the dominant role of eye movements and how they link perception and action. Either way, regardless of the mechanism underlying these eye movements, the paradigm used here offers a useful approach to directly readout dynamical internal beliefs in real-time, simply by tracking subjects’ eyes.

## METHODS

### Experimental Model and Subject Details

Three rhesus macaques (all male, 7-8 yrs. old) and ten human subjects (six males, all adults in the age group 18-32 yrs.) participated in the experiments. All but one subject were unaware of the purpose of the study. All surgeries and experimental procedures were approved by the Institutional Review Board at Baylor College of Medicine, and were in accordance with National Institutes of Health guidelines. All human subjects signed an approved consent form. In the following sections, the term subject is used to denote both monkey and human subjects, unless specified otherwise or implied by the context.

### Method Details

#### Experimental setup

Monkeys were chronically implanted with a lightweight polyacetal ring for head restraint, and scleral coils for monitoring eye movements (CNC Engineering, Seattle WA, USA). At the beginning of each experimental session, monkeys were head-fixed and secured in a primate chair placed on top of a platform (Kollmorgen, Radford, VA, USA). A 3-chip DLP projector (Christie Digital Mirage 2000, Cypress, CA, USA) was mounted on top of the platform and rear-projected images onto a 60 x 60 cm tangent screen that was attached to the front of the field coil frame, ~30cm in front of the monkey. The projector was capable of rendering stereoscopic images generated by an OpenGL accelerator board (Nvidia Quadro FX 3000G).

Human subjects wore a custom-fit thermoplastic mask (CIVCO Medical Solutions) that was screwed to the back of the chair to restrain their head. The mask was mounted with a binocular eye tracker (ISCAN Inc.) to record the position of the subjects’ pupils at 60Hz. All other aspects of the setup were similar to the one used for monkeys, but with subjects seated 67.5cm in front of a 149 × 127 cm^2^ (width × height) rectangular screen. Although humans and monkeys were head-fixed, they were both free to move their eyes when performing the task, except under one experimental manipulation in humans (noted towards the end of the section below).

#### Behavioural Task

Subjects used an analog joystick (M20U9T-N82, CTI electronics) with two degrees of freedom and a circular displacement boundary to control their linear and angular speeds in a virtual environment. This virtual world comprised a ground plane whose textural elements had limited lifetime (~250ms) to avoid serving as landmarks. The ground plane was circular with a radius of 70m (near and far clipping planes at 5cm and 4000cm respectively), with the subject positioned at its center at the beginning of each trial. Each texture element was an isosceles triangle (base × height: 8.5 × 18.5 cm^2^) that was randomly repositioned and reoriented anywhere in the arena at the end of its lifetime, making it impossible to use as a landmark. The maximum linear and angular speeds were fixed to *v*_max_ = 2ms^−1^ and *ω*_max_ = 90°/s respectively, and the density of the ground plane was either held fixed at *ρ* = 2.5 elements/m^2^ or varied randomly between two values (*ρ* = 2.5 elements/m^2^ and *ρ* = 0.1 elements/m^2^) in a subset of recording sessions (see below). The stimulus was rendered as a red-green anaglyph and projected onto the screen in front of the subject’s eyes. Subjects wore goggles fitted with Kodak Wratten filters (red #29 and green #61) to view the stimulus. The binocular crosstalk for the green and red channels was 1.7% and 2.3% respectively.

Human subjects pressed a button on the joystick to initiate each trial, and the task was to steer to a random target location that was cued briefly at the beginning of the trial (Fig 1A). Monkeys performed the same task, but each trial was programmed to start after a variable random delay (0.5 – 1.1s) following the end of the previous trial. The target was a circular disc of radius 20cm whose luminance was matched to the texture elements. It appeared at a random location between θ = ±40° of visual angle at a distance of *r* = 0.7 − 4m (up to 6m for human subjects) relative the subject at the beginning of the trial. For human subjects, the target disappeared after one second, which was a cue for the subject to start steering, and the joystick controller was activated. In the case of monkeys, the target only appeared on the screen for 300ms, and the joystick was always active.

Monkeys typically performed two blocks of ~750 trials in each experimental session, and received feedback at the end of each trial. Monkeys performed a total of ~6,000 trials (4 sessions) each. Eye tracking was performed either using scleral coils (monkey Q & B) or a head-mounted eye tracker (monkey S). In one of the above recording sessions in each monkey, we saved the stimulus movie and replayed them to the animal at the end of the block. Both the visual stimulus and the schedule of rewards during this replay block were identical to the active navigation block, with the only difference being that the joystick was withheld and monkeys passively viewed the stimulus. Furthermore, a subset of the recording sessions (two sessions in each monkey) contained two randomly interleaved sets of trials that differed in terms of the density of optic flow (*ρ* = 0.1 elements/m^2^ and *ρ* = 2.5 elements/m^2^).

Of the ten human subjects, five subjects performed a total of 600 trials spread equally across three blocks. The blocks were identical in all respects, except no feedback was provided at the end of the trials in the first and third blocks. The purpose of using this block structure was to study how feedback affected learning in humans. Although data collected in the absence of feedback (first and last blocks) are briefly described in **Fig. S1**, the key results of the paper are based only on data collected during the intermittent block with feedback. Furthermore, during the block with feedback, the performance of human subjects typically stabilized within fifty trials (**Fig. S1B**). Because we wanted to ensure that the performance was stable during the course of testing, we ignored the first fifty trials of this block for all our analysis (Figs 1,2,4). The remaining five human subjects participated in a version of the experiment that was designed to study the effect of inhibiting eye movements on task performance (Fig 6). These subjects first performed a block of fifty trials with feedback to allow their performance to stabilize. Following this pre-training block, they performed a test block comprising 400 trials of a version of this task in which a fixation cross was overlaid on top of the target in each trial, again with feedback. In a random subset of trials (50%), this fixation cross remained on the screen even after the target disappeared and subjects were instructed to maintain fixation on the cross while steering to the target. The location of the cross remained fixed in screen coordinates and thus carried no dynamic information about stimulus location.

#### Feedback

Monkeys received binary feedback at the end of each trial. They received a drop of juice if, after stopping, they were within 0.6m away from the center of the target. No juice was provided otherwise. The fixed reward boundary of 0.6m was determined using a staircase procedure prior to the experiment to ensure that monkeys received reward in approximately two-thirds of the trials.

Human subjects received a somewhat richer, adaptive feedback in the form of a bullseye pattern that appeared on the ground at the end of steering. The bullseye was centered on the target, with the innermost region having the highest luminance. The pattern comprised of five zones (**Fig S1A**), and the radii of the rings were continuously scaled (up or down by 5%) during the experiment using a 1-up 2-down staircase procedure. Additionally, an arrowhead pointing to the target also appeared on the ground in front of the subjects, colored green or red depending on whether the subject’s stopping position was inside or outside the reward boundary. The adaptive feedback procedure ensured that human subjects, like monkeys, stopped within the reward boundary in roughly two-thirds of the trials. Unlike monkeys, human subjects did not receive juice at the end of each successful trial, but instead received monetary compensation that was commensurate with their performance.

#### Stimulus and Data acquisition

All stimuli were generated and rendered using C++ Open Graphics Library (OpenGL) by continuously repositioning the camera based on joystick inputs to update the visual scene at 60 Hz. The camera was positioned at a height of 1m above the ground plane (10cm for monkeys). Spike2 software (Power 1401 MkII data acquisition system from Cambridge Electronic Design Ltd.) was used to record and store the target location (*r*^*^, *θ*^*^), subject’s position (*r*, *θ*), horizontal positions of left and right eyes (*α*_*l*_ and *α*_*r*_), vertical eye positions (*β*_*l*_ and *β*_*r*_) and all event markers for offline analysis at a sampling rate of 833 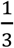 Hz.

#### Model predicted eye position

To test whether subjects’ eyes tracked the location of the (invisible) target, we generated predictions for subjects’ instantaneous eye positions by assuming that they maintained fixation at the center of the target. (*x*_*t*_, *y*_*t*_, *z*_*t*_) denotes the location of the target relative to the mid-point of the subject’s eyes at time *t*. The *mean* predicted lateral displacement (relative to fixating at the point (0, ∞, 0)) of the left and right eyes 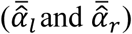 are geometrically related to the target location and the inter-ocular distance (2∆) as (**Figs S4B**):

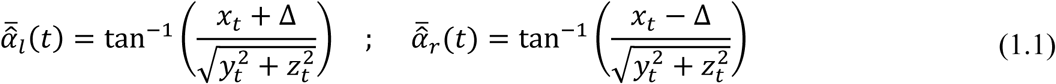

Likewise, the vertical displacement of the two eyes 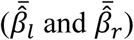 should be:

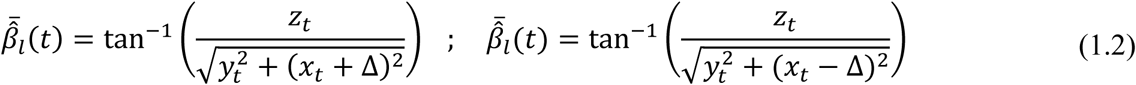

Note that *z*_*t*_ is determined entirely by the camera height and hence time-invariant. In contrast, *x*_*t*_ and *y*_*t*_ change continuously as the subject steers to the target, and are both equal to zero in the special case when the subject’s location coincides with the center of the target. The predicted eye positions also have *variances* associated with them, which we derive in a later section (equation 4).

### Quantification and Statistical Analysis

Customised MATLAB code was written to analyse data and to fit models. Depending on the quantity estimated, we report statistical dispersions either using 95% confidence interval, standard deviation, or standard error in the mean. The specific dispersion measure is identified in the portion of the text accompanying the estimates. For error bars in figures, we provide this information in the caption of the corresponding figure. We report exact *p*-values for all statistical tests, and describe the outcome as significant if *p* < 0.05.

#### Bias estimation

We regressed (with an intercept term) each subject’s response positions (*r*, θ) against target positions (*r*^*^, θ^*^) separately for the radial (*r* vs *r*^*^) and angular (θ vs θ^*^) co-ordinates, and the radial and angular multiplicative biases were quantified as the slope of the respective regressions (Fig 1F). The intercept terms of the regression models denote additive bias. For each subject, we estimated the 95% confidence intervals for the biases by bootstrapping.

#### Psychometric analysis

As described in the section on feedback, reward boundaries were chosen to ensure that all subjects correctly stopped within the reward zone in about two-thirds of the trials. However, the precise radius of these boundaries varied across human subjects, as well as between humans and monkeys. To objectively compare the performance of different subjects on a common scale, we performed ROC analysis as follows. For each subject, we first constructed a psychometric function by calculating the proportion of correct trials as a function of (hypothetical) reward boundary (Fig 1G). In keeping with the range of target distances used for the two species, we varied the reward boundary between 0–4m for monkeys and 0–6m for human subjects. Whereas an infinitesimally small boundary will result in all trials being classified as incorrect, a large enough reward boundary will yield near-perfect accuracy. To define a chance-level psychometric function, we repeated the above procedure but now by shuffling the target locations across trials, thereby destroying the relationship between target and response locations. Finally, we obtained the ROC curve by plotting the proportion of correct trials in the original dataset (true positives) against the shuffled dataset (false positives) for each value of hypothetical reward boundary. We used the area under this ROC curve to obtain an accuracy measure that was independent of the reward boundary used for various subject.

#### Characterizing eye position

For convenience, we express the subject’s *actual* eye position using the following three standard degrees of freedom: (*i*) Conjunctive horizontal movement of the two eyes or ‘lateral version’ quantified here as the mean lateral position of the two eyes, α = (*α*_*l*_ + *α*_*r*_)/2, (*ii*) Conjunctive vertical movement of the two eyes or ‘elevation’ quantified here as *β* = (*β*_*l*_ + *β*_*r*_)/2, (*iii*) Disjunctive horizontal eye movements or ‘vergence’ quantified here as *γ* = (*α*_*l*_ − *α*_*r*_)/2. Disjunctive eye movements along the vertical direction (vertical vergence) were an order of magnitude smaller than the precision of our measurements, and therefore we ignore them in all our analyses. We also transformed the *predicted* eye positions given by Equation 1 into the above three degrees of freedom using analogous definitions to obtain 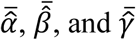.

#### Saccade detection and pre-processing

We estimated the instantaneous speed of eye movements as 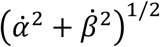 where *α* and *β* denote lateral version and elevation respectively (as defined above), and a dot denotes a time derivative. Saccades were detected by identifying the time points at which the speed of eye movements crossed a threshold of 200º/s from below (a threshold of 50º/s yielded similar results). Although saccades were mostly confined to periods immediately following target onset and end of steering (Fig 2B), we removed a period of 100ms immediately following the onset of saccades for visualizing the time-course of eye movements during the trial (Fig 2E) and for all subsequent temporal analyses described below. We verified that this procedure had minimal effect on the results. In approximately 10% of the trials in monkeys and ~30% in human subjects, the subject travelled beyond the target. The predicted eye positions towards the end of these trials were outside the range that was physically possible. Therefore, we removed time points at which any of the four predicted components of eye movements in Equation 1 exceeded 60º before further analysis. Such time points constituted less than 3% of the dataset, and including them did not qualitatively alter the results.

#### Comparing predicted and observed eye positions

Let **φ**_*t*_ = (*α*_*t*_, *β*_*t*_) and 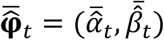 denote the observed and mean predicted eye positions respectively at time *t*. For each subject, we computed the square root of the fraction of variance in their eye movements explained by the predictions:

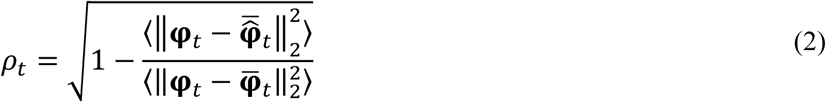

where ║∙║_2_ denotes the *L*_2_ norm, 〈∙〉 denotes expectation across trials, and 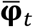 denotes the mean observed eye position across trials at time *t*. Because the predictions are based on a model that assumes subjects’ eyes track the center of the target, we call *ρ* the ‘*target*-*tracking index*’, or simply ‘*tracking-index*’. A value of 1 corresponds to perfect prediction while zero implies that the predictions were no better than the mean observation. In principle, the deviation from the predictions can be larger than the intrinsic variability of the data. We clipped the target-tracking index to zero whenever this happened. Since trial durations were variable, we aligned all trials relative to the time at which the target was turned off (*t* = 0) to estimate the time course of tracking coefficient 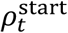 ∀ *t* ∈ [0, 1.8*s*]. 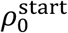 corresponds to the similarity between observed and predicted eye position at the moment when the target was turned off (Fig 2F). We also computed the tracking coefficient by aligning trials with respect to the end of steering (*t* = *T*) to estimate 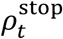 ∀ *t* ∈ [−1.2s, 0]. To visualize the time-course of the tracking coefficient, we plot both 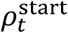 and 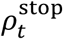 with a break in the *x*-axis (Fig 2G, 4 & 5). To assess standard errors and statistical significance of differences between tracking coefficients from pairs of conditions (e.g. rewarded vs unrewarded trials), we used a bootstrap test with 10,000 bootstrap samples.

#### Correlation between saccade amplitude and target-tracking error

Because saccades were not very frequent while steering, we pooled data from all subjects (separately for monkeys and humans) for analyzing saccades. The amplitude of saccades was taken to be the average displacement of the position of the two eyes from saccade onset to 100ms later (∆**φ** = (∆*α*^2^ + ∆*β*^2^)^1/2^; Fig 3B). We quantified the effect of saccadic eye movements on target-tracking error by computing the saccade-triggered average (STA) of the tracking error within a 400ms window centered around time *t*_*s*_ of saccade onset (*i.e*., 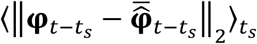 ∀ *t* ∈ [−0.2*s*, 0.2*s*]). To quantify the precise relationship between saccade amplitude and tracking error, we simultaneously regressed horizontal and vertical amplitudes of the saccade (∆*α* and ∆*β*) on horizontal and vertical tracking errors 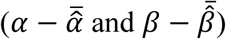, respectively, at various lags between ±0.5*s* with *l*^2^ regularization.

#### Estimation of position uncertainty

We estimated subjects’ position uncertainty by binning the 2D space into 10×10 cm^2^ bins. For each bin, we computed the variance in the subject’s stopping position across trials in which targets fell in that bin. The resulting spatial map of variability was then convolved with a two-dimensional isotropic Gaussian kernel of width 40cm (equal to the diameter of the target) to yield a smooth estimate of variability as a function of space (Fig 4B – *left*). Because subjects aimed to stop on the target, variability in their stopping position can be interpreted as the uncertainty in subjects’ posterior estimate about their own position.

#### Deriving an upper bound on the target-tracking index

Once the target disappears, subjects no longer get to directly observe it. To reach the target location, they update their beliefs about the relative location of the target by integrating their self-motion, which in turn must be estimated from the observed optic flow. Even if those beliefs are accurate on average, the uncertainty in believed target location will grow over time on any given trial due to noise both in the observations and in the integration process. Consequently, the degree to which subjects’ eyes can track the target (quantified by the tracking index, *ρ*) should decrease over time. Using the variability in subjects’ stopping positions to model their uncertainty in their believed location (see section above), we derived an approximate upper-bound on the temporal dynamics of the tracking-index *ρ*_*t*_ at time *t* assuming inter-ocular distance ∆≈ 0:

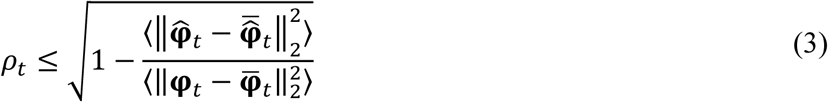

where 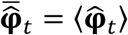 denotes the mean predicted eye position at time *t*. Note that this represents an upper-bound insofar as the variability in subject’s stopping positions stems entirely from uncertainty in their believed location. To derive this approximate bound, we first used the first-order Taylor series approximation of equation (1) to express the variance of the predicted eye position 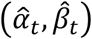 in terms of the variance of the relative target position (*x*_*t*_, *y*_*t*_, *z*_*t*_) as: Var(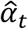) = (*∂f*/*∂x*)^2^ Var(*x*_*t*_) + (*∂f*/*∂y*)^2^ Var(*y*_*t*_)and Var(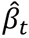) = (*∂g*/*∂x*)^2^ Var(*x*_t_) + (*∂g*/*∂y*)^2^ Var(*y*_*t*_), where *f*(*x*_*t*_, *y*_*t*_, *z*_*t*_) = tan^−1^ (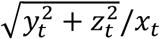) and *g*(*x*_*t*_, *y*_*t*_, *z*_*t*_) = tan^−1^ (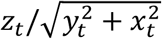) from equation (1), and we have used the fact that Var(*z*_*t*_) = 0 because there is no motion component perpendicular to the ground plane. Substituting the derivatives, we get:

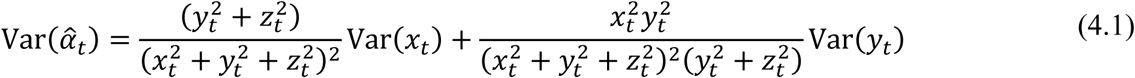

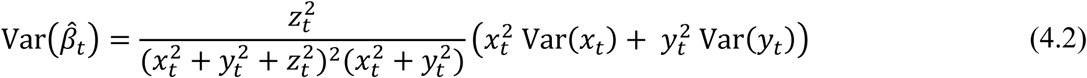

The above equations are based on first-order Taylor series approximation and hold as long as the higher-order terms are relatively small (**Fig S9**). Although we cannot not directly measure Var(*x*_*t*_) and Var(*y*_*t*_), we could estimate them from the data (see previous section) and use it to determine the variability in predicted eye positions given by **equation (4)**. Variability in the predictions then implies a lower bound in the mean squared error achievable by any observation φ_*t*_: 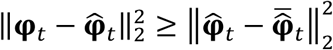. Substituting this in **(2)**, we obtain an upper bound on the tracking-index given by **equation (3)**. Note that, in deriving this approximate upper-bound, we ignored the noise in generating an eye movement to an intended location (process noise). So in principle, it is possible to derive a tighter bound by incorporating it. Note that as subjects approach the target, *x*_*t*_ and *y*_*t*_ approach zero, whereas the uncertainty grows so both Var(*x*_*t*_) and Var(*y*_*t*_) increase. Together, this leads to an increase in the variance of the predicted eye positions (equation 4) and consequently, a gradual decrease in the fraction of explainable variance over time (equation 3).

#### Comparing behavioural and target-tracking errors

To test whether poor target-tracking was associated with poor behavioural accuracy, we estimated the correlation between behavioural and target-tracking errors across trials of individual recording sessions. Behavioural error was given by the Euclidean distance between the target location and the subject’s response (stopping location) on individual trials, while the target-tracking error was given by the Euclidean distance between actual and predicted eye position, averaged over the entire time period of the trial, except for the last 300ms (as the predictions typically broke down when the subject was too close to the target). Because trial difficulty could affect both errors thereby inducing spurious correlations, we estimated the null distribution of correlations using a shuffling procedure where we grouped the trials from each recording session into ten quantiles based on target distance and shuffling only trials within the same group. The results were quite robust to the number of quantiles.

#### Assessing performance in the fixation task

To assess the behavioural effect of inhibiting eye movements, we compared human subjects’ performance across ‘eyes-moving’ and ‘eyes-fixed’ trials. Because we did not control for fixation breaks that happened in the ‘eyes-fixed’ condition during the experiment, we identified and removed such trials offline. Specifically, we removed the trials in which the temporal standard deviation (*σ*) of subject’s eye position during the trial (*i.e.* from the time when the target disappeared until the end of steering) exceeded 3° (roughly half-width of the fixation cross), from our analysis (~10% of the fixation trials across all subjects). The standard deviation was quantified as *σ* = 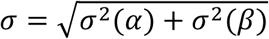 where *σ*(*α*) and *σ*(*β*) denote the temporal standard deviation of lateral version and elevation respectively. To evaluate the role of eye movements, we compared subjects’ performance in the fixation trials (‘eyes-fixed’) with trials that did not require fixation (‘eyes-moving’). For both sets of trials, we computed ROC curves for distinguishing ‘rewarded’ and ‘unrewarded’ trials (see section ‘psychometric analysis’ above) and used a paired *t*-test to test whether the mean area under the curves were different. We also computed the correlation between target and response locations and then used a paired *t*-test to test whether there was a significant difference between the correlation coefficients in the two sets of trials across subjects (Fig 6D).

## ACKNOWLEDGEMENTS

We thank Samir Saidi for assisting with the human experiments, Jing Lin and Jian Chen for their help in programming the stimulus. This work was supported by grants EY025538 and R01-DC014678. G.C.D. was supported by EY016178.

## AUTHOR CONTRIBUTIONS

Conceptualization, K.J.L., G.C.D., X.P. and D.E.A.; Methodology, K.J.L., E.A., X.P. and D.E.A., Investigation, K.J.L., E.A. and E.N.; Data Curation, K.J.L. and E.A.; Formal Analysis, K.J.L.; Writing – Original draft, K.J.L.; Writing – Review & Editing, K.J.L., E.A., G.C.D., X.P. and D.E.A.; Funding Acquisition – G.C.D., X.P. and D.E.A.; Supervision, X.P. and D.E.A.

