## Supplementary figures for "Tracking the mind’s eye: Primate gaze behavior during virtual visuomotor navigation reflects belief dynamics"

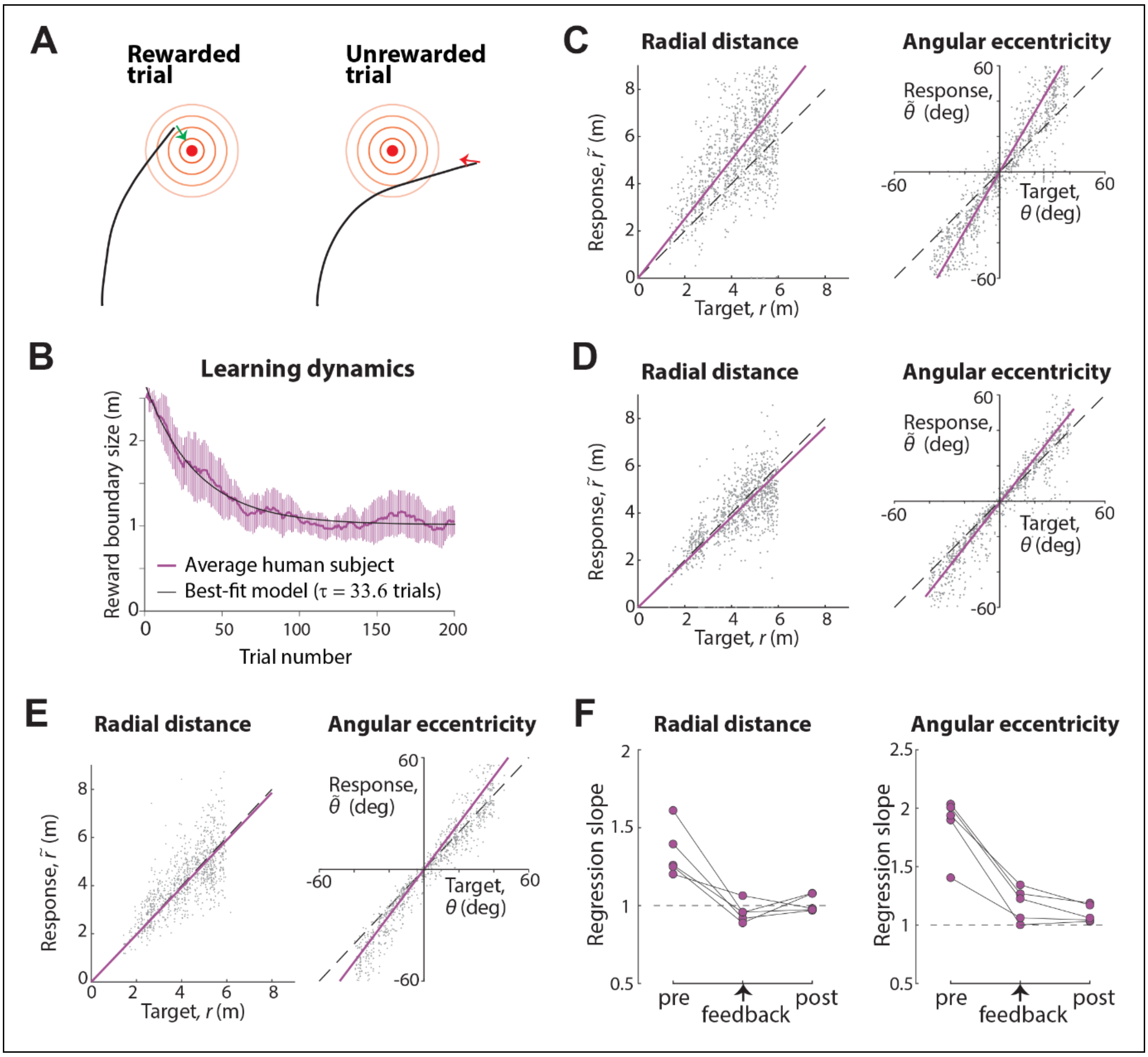


**Figure S1. Behavioural performance of human subjects. A. Feedback for humans.** At the end of the movement, a bullseye pattern (concentric rings) centred on the target appeared on the ground plane to indicate the magnitude of error. The arrowhead served both as binary feedback (green – correct, red – incorrect) and to indicate the direction of error. **B. Learning rate.** The radius of the bullseye pattern was adaptively scaled using a staircase procedure (see **Methods**). Purple curve shows the average radius ($R$) across subjects as a function of trial number. Error bars denote ±1 standard deviation. The black curve shows the exponential function $R(n) \propto \exp(-n/\tau)$ fit to data. **C. Pre-feedback performance.** *Left*: Comparison of the radial distance of the response (final stopping position) against radial distance of the target across all trials of all human subjects during the first experimental block (pre-feedback block). *Right*: Angular eccentricity of subjects' response vs target angle. Black dashed lines have unity slope (unbiased performance). **D. Performance improved with feedback.** Similar to (C), but for the last 150 trials of the block with feedback (second block). **E. Performance improvement persisted after removing feedback.** Similar to (C), but for the final block of trials with feedback withheld (third block). **F.** Regression slopes of individual subjects across all three blocks (pre-feedback, feedback and post-feedback).

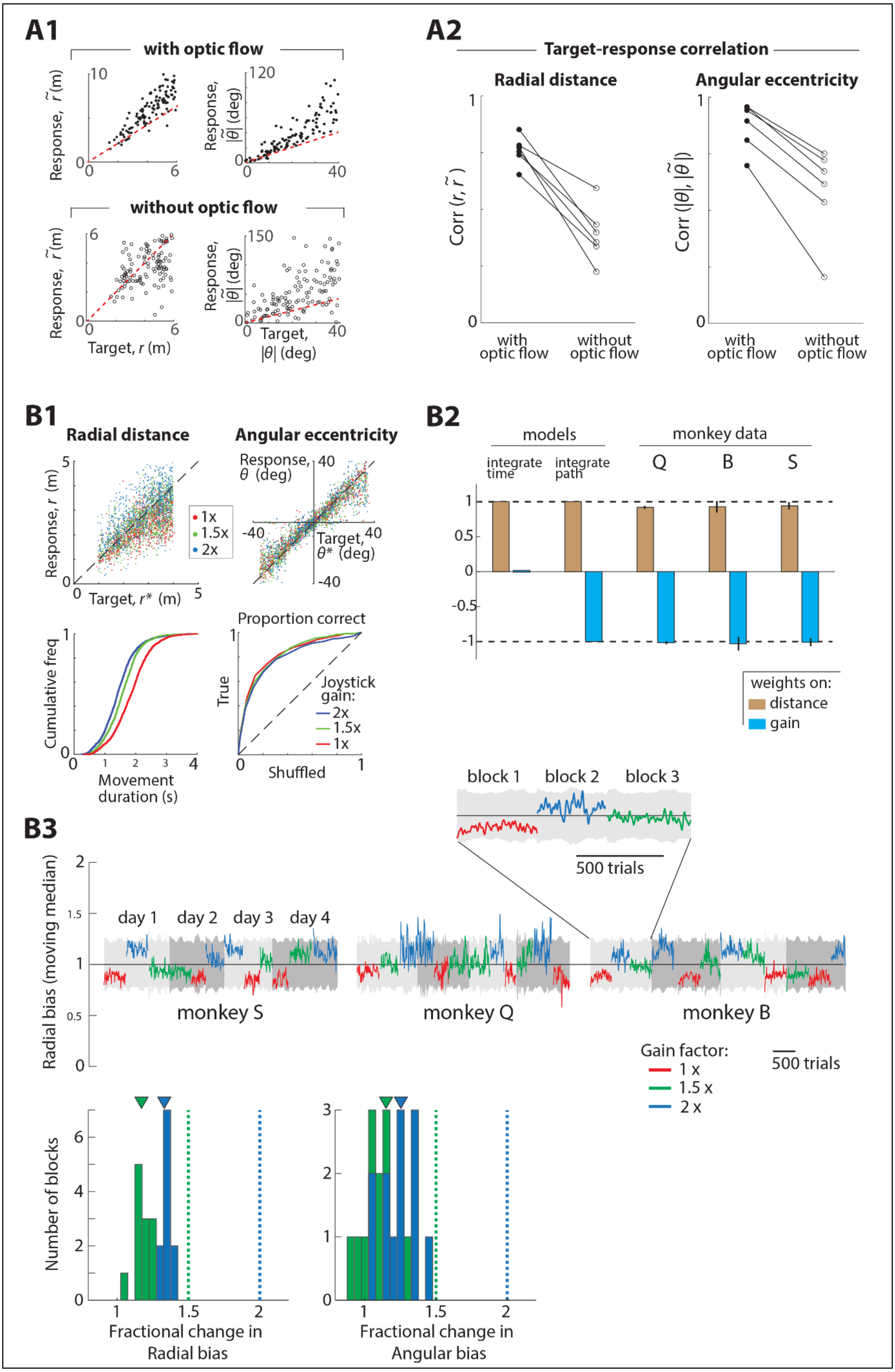
A1
with optic flow
Response, r̃ (m)
10
0
0
6
Response, |θ̃| (deg)
120
0
0
40
without optic flow
Response, r̃ (m)
6
0
0
6
Target, r (m)
Response, |θ̃| (deg)
150
0
0
Target, |θ| (deg)
40
A2
Target-response correlation
Radial distance
Angular eccentricity
Corr (r, r̃)
Corr (|θ|, |θ̃|)
1
0
with optic flow
without optic flow
B1
Radial distance
Angular eccentricity
Response, r (m)
5
0
0
Target, r* (m)
5
1x
1.5x
2x
Response, θ (deg)
40
-40
Target, θ* (deg)
40
-40
Cumulative freq
1
0
0
1
2
3
4
Movement duration (s)
Proportion correct
True
1
0
Shuffled
1
Joystick gain:
2x
1.5x
1x
B2
models
monkey data
integrate time
integrate path
Q
B
S
1
0.5
0
-0.5
-1
weights on:
distance
gain
B3
block 1
block 2
block 3
500 trials
Radial bias (moving median)
2
1.5
1
0.5
0
day 1
day 2
day 3
day 4
monkey S
monkey Q
monkey B
Gain factor:
1 x
1.5 x
2 x
500 trials
Number of blocks
6
4
2
0
1
1.5
2
Fractional change in Radial bias
3
2
1
0
1
1.5
2
Fractional change in Angular bias

**Figure S2. Subjects rely on optic flow to perform the task. A. Human subjects.** [*Data from Lakshminarasimhan et al. (2018)*] Texture elements constituting the ground plane were removed in a random subset of trials to block optic flow cues. Subjects did not receive feedback at the end of the trials. (**A1**) Radial and angular response of an example human subject with (top panels) and without (bottom panels) optic flow cues. The subject's overall variability was much larger without optic flow cues. (**A2**) Across subjects, removing optic flow cues induced a significant decrease of target-response correlations in both radial distance [Corr($r$,$\tilde{r}$) : 0.76±0.06 with optic flow, 0.39±0.12 without optic flow, $p$=4.1×10$^{-4}$, paired $t$-test] and angle [Corr($|\theta|$,$|\tilde{\theta}|$) : 0.88±0.1 with optic flow, 0.58±0.2 without optic flow, $p$=4.8×10$^{-4}$] suggesting that subjects relied heavily on optic flow cues. **B. Monkeys.** In separate blocks, we manipulated the gain of the joystick controller to alter the sensorimotor mapping learned by the monkey (gain values of 1x (baseline), 1.5x and 2x). Each block comprised around 500 trials and the order of the blocks were randomized across days. (**B1**) *Top panels*: Radial (*left*) and angular (*right*) response of an example monkey during trials from different gain conditions (red: 1x, green: 1.5x, blue:2x) showing that stopping positions were close to target locations under all conditions. *Bottom left*: Cumulative distribution of travel times for the different gain conditions. The mean travel time significantly decreased when gain was increased (mean ± std for 1x gain: 1.4$\pm$0.1s, 2x gain: 2.1$\pm$0.2s; $p$ < 10$^{-5}$, paired $t$-test) implying that monkeys adapted to the different gain values by adjusting their travel duration appropriately. *Bottom right*: Average ROC curves of the monkeys, obtained by plotting their true proportion of correct trials (from unshuffled data) against the corresponding chance-level proportions (from shuffled data) for a range of reward windows separately for each gain condition. The area under the curve (AUC) was comparable under all three conditions (mean AUC ± std. for 1x gain: 0.82$\pm$0.03, 1.5x gain: 0.85$\pm$0.04, 2x gain: 0.79$\pm$0.06) implying that their task performance was not affected by manipulating the joystick gain . (**B2**) We simultaneously regressed the travel time against initial target distance ($r$) and joystick gain ($g$) in the log space across trials [$\log(T) = w_r \log(r) + w_g \log(g)$]. Travel time of an ideal path integrator would depend on changes in gain and have negative regression weight on the gain term ($w_g = -1$) whereas travel time for pure time integration or dead-reckoning would be insensitive to gain changes ($w_g = 0$). Across the set of all rewarded trials, the regression weight $w_g$ on joystick gain was not significantly different from $-1$ in all three monkeys (95% confidence interval (CI) of regression weight, Monkey B: $[-0.99, -1.04]$ , Monkey S: $[-0.95, -1.17]$, Monkey Q: $[-0.96, -1.09]$). (**B3**). *Top panels*: Time-course of the radial bias (explained below) over the series of trials from three different blocks of the experiment (each block had a different gain; *red:* 1x, *green:* 1.5x, *blue:* 2x) from four consecutive days of each monkey. Day 1 of each monkey corresponds to their first exposure of control gain manipulation. Single-trial radial bias was computed by first taking the ratio of stopping distance and target distance on each trial, followed by a moving median with a window of twenty-five trials (window truncated as required for trials in the beginning and end of each block). Inset shows results from day1 of monkey B. Regression analysis on these time-courses revealed that there was neither gradual adaptation across trials within each block, not across different days suggesting that the adaptation was rapid and therefore largely mediated by optic flow cues (95% CI of regression weight on trial number: 1.5x gain: $[-0.00002, 0.0001]$, 2x gain: $[-0.0003, 0.001]$; regression weight on day number: 1.5x gain: $[-0.1, 0.05]$ , 2x gain: $[-0.12, 0.06]$). Grey shaded regions show the running median of the bounds on radial bias within which monkeys get reward. *Bottom left panel*: The histogram of mean ratios of radial bias in the block with a gain factor of 1.5x and the block with gain factor of 1x is shown in green (comparison of 2x to 1x in blue). *Bottom right*: Similar to bottom left, but showing ratios of angular bias. The median ratios were significantly smaller than expected by lack of adaptation (median ratio $\pm$ IQR of radial bias: 1.5x vs 1x: 1.18$\pm$0.05, 2x vs 1x: 1.33$\pm$0.05; median ratio $\pm$ IQR of angular bias: 1.5x vs 1x: 1.19$\pm$0.12, 2x vs 1x: 1.24$\pm$0.2).

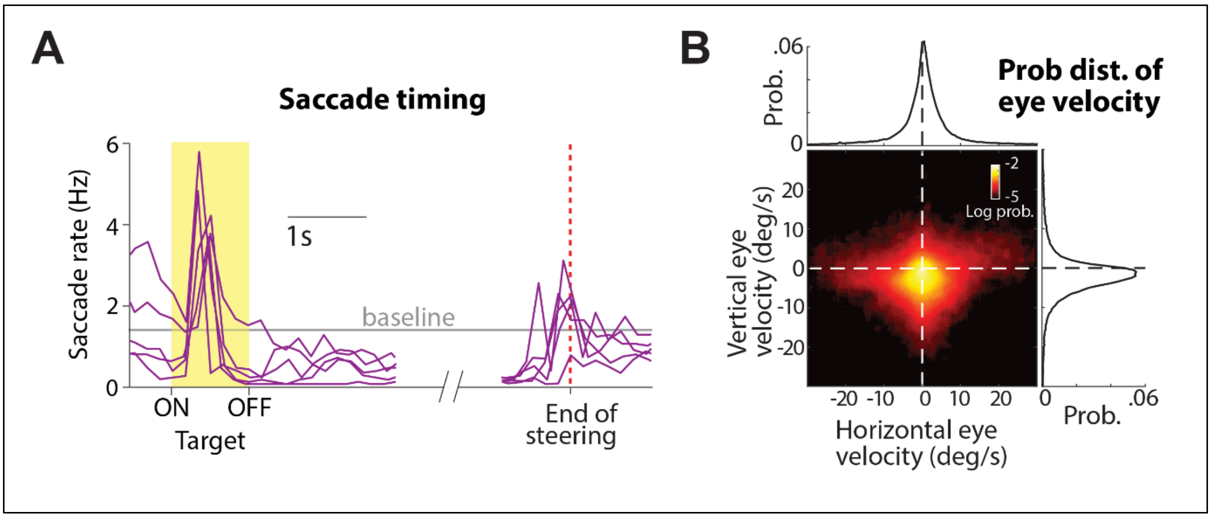


**Figure S3. Saccade timing and eye velocity of human subjects. A.** The trial-averaged saccade rate of individual human subjects. Trials were aligned relative to target presentation (*shaded yellow* to the left of the break on the x-axis) and end of movement (*red dashed line* after the break). Note that targets were visible for a period of one second. **B.** The joint probability density of distribution over horizontal and vertical eye velocities, averaged across human subjects, while they steered towards the target.

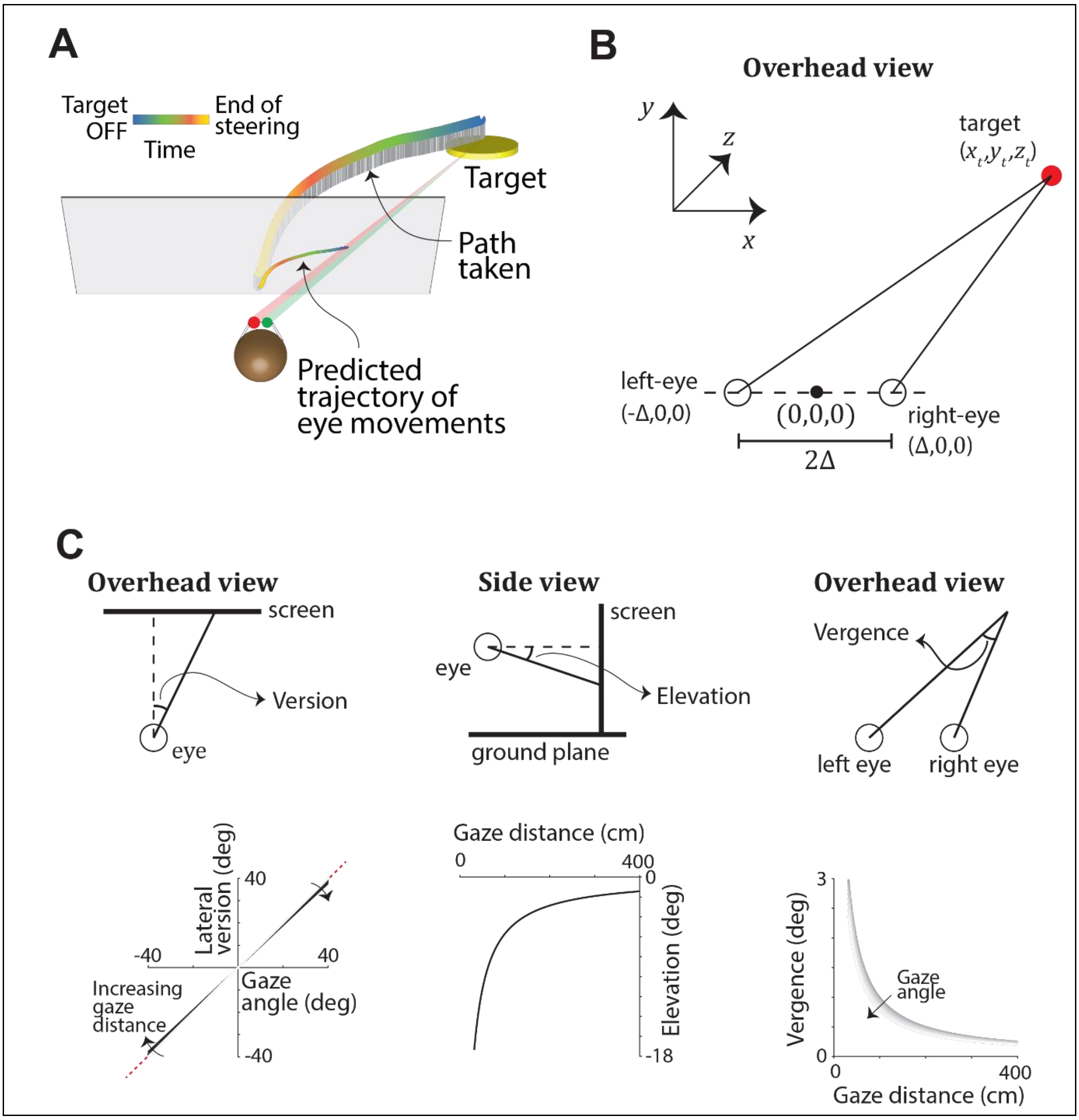


**Figure S4. Modelling angular eye position from gaze location. A.** Graphical illustration of the expected dynamics of subject's eye movements (average of the two eyes, projected onto the plane of the screen) while steering to the target. Time is coded by colour. **B.** The instantaneous three-dimensional egocentric position of the target (red) is used to generate theoretical predictions for the subject's eye position, assuming they maintained fixation at the centre of the target throughout the trial (**Methods – Equation 1**). **C.** *Top*: Eye position is characterized using three degrees of freedom: *Lateral version*, which measures the average deviation of the two eyes from the sagittal plane (*left*), *Elevation*, which measures the average deviation of the eyes from the transverse plane (*middle*), and *Vergence* which measures the difference between the lateral position of the two eyes (*right*, see **Methods** for quantitative definitions). *Bottom*: Theoretical dependence of the magnitudes of lateral version, elevation, and vergence on gaze distance and gaze angle (inter-ocular distance, $\Delta$ = 3.5cm; eye height, $z$ = 10cm).

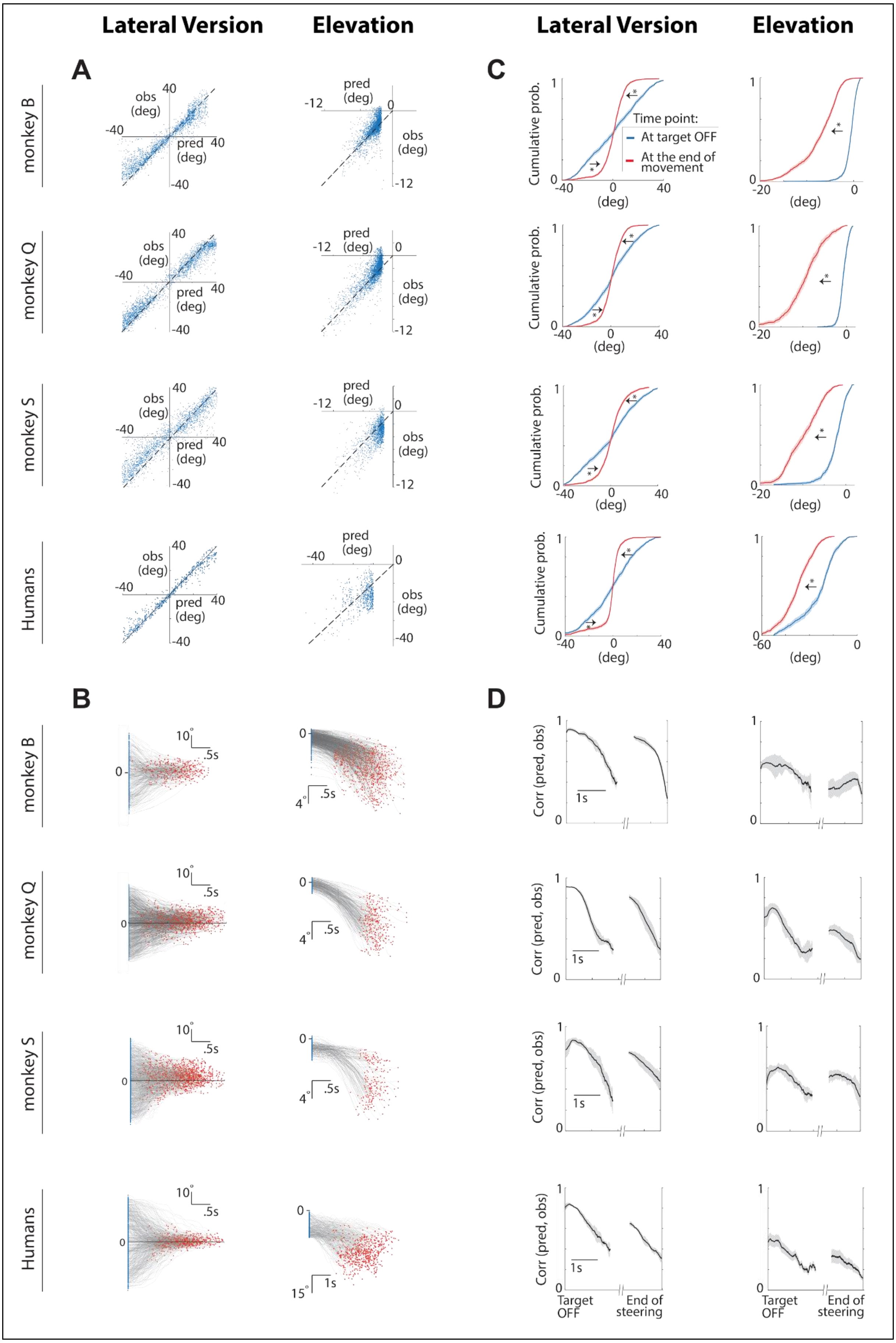
Lateral Version
Elevation
Lateral Version
Elevation
A
B
C
D
monkey B
monkey Q
monkey S
Humans
obs (deg)
pred (deg)
Cumulative prob.
Time point:
At target OFF
At the end of movement
(deg)
Corr (pred, obs)
1s
Target OFF
End of steering

**Figure S5. Temporal dynamics of lateral version and elevation. A.** Comparison of the two components (lateral version - *left*, elevation – *right*) of predicted and true eye positions in a random subset of trials sampled from individual monkeys (rows 1-3) and all human subjects (combined, last row) at the moment when the target is turned OFF. **B.** The time-course of the above two components of eye movements during a random subset of trials from monkeys and humans. Blue and red dots denote the times at which the target was turned OFF and the end of movement, respectively. **C.** The empirically estimated cumulative density functions of the distribution over the subjects' lateral version (*left*) & elevation (*right*) at the moment when the target was turned OFF (*blue*) and at the end of movement (*red*). Shaded regions denote standard errors obtained by bootstrapping. For each of the two components, we used a bootstrap test with 10,000 bootstrap samples to determine whether parameters of the distributions estimated at the two time points were different. Specifically, we qualitatively expect the *magnitude* of elevation to increase over time (**Fig S3**), so we tested whether the magnitude of the median was significantly greater at the end of movement. On the other hand, we expect the magnitude of lateral version to approach zero over time (**Fig S3**), so we tested whether the inter-quartile range (a measure of dispersion) was significantly smaller at the end of movement. Asterisks denote significant difference ($p<0.001$) and n.s. denotes not significant according to the bootstrap test described above (*i.e.* whenever $p >= 0.001$). **D.** Time-course of the Pearson's correlation between the predicted and observed values of version (*left*) and elevation (*right*) computed by aligning trials with respect to both the time when the target was turned OFF (denoted by the y-axis on the left) as well as the end of movement (denoted by the y-axis on the right). For humans, we pooled trials from all five subjects for the analysis shown here.

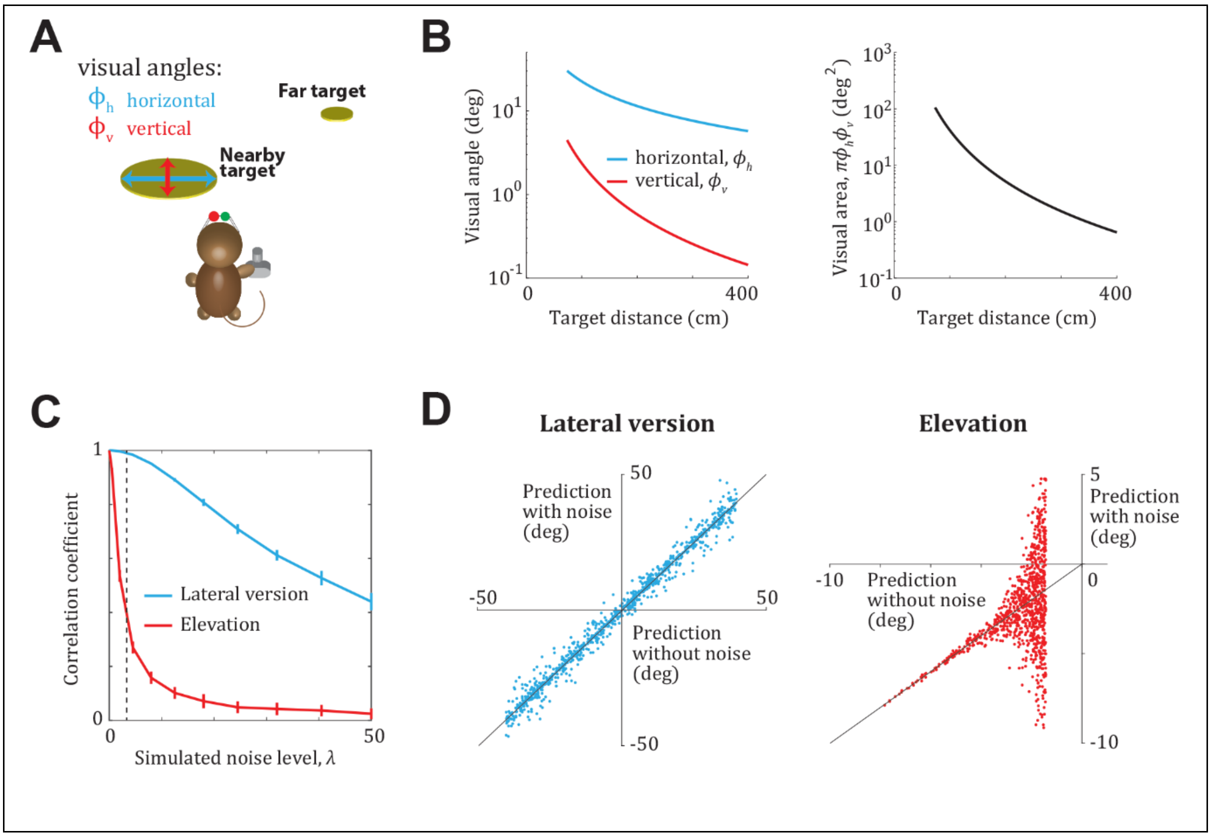


**Figure S6. Modelling the effect of target distance on eye movement variability. A.** Graphical illustration showing the difference in visual size of example nearby and distant targets. **B.** *Left*: Theoretical values of vertical ($\varphi_v$, red) and horizontal ($\varphi_h$, blue) visual angles subtended by the target, as a function of target distance (calculated using **Methods – Equation 1** for a circular target of radius 20cm). *Right*: Compared to the nearest targets, visual area ($\pi\varphi_h\varphi_v$) drops by roughly two orders of magnitude for the farthest targets. **C.** Assuming the noise is inversely proportional to visual area of the target, we numerically computed the correlation coefficients between noisy and noiseless predictions, as a function of noise by varying the constant of proportionality $\lambda$. **D.** Effect of noise on lateral version and elevation for $\lambda = 3$ (corresponding to the dashed vertical line in C). Model parameters: inter-ocular distance = 3.5cm; eye height = 10cm.

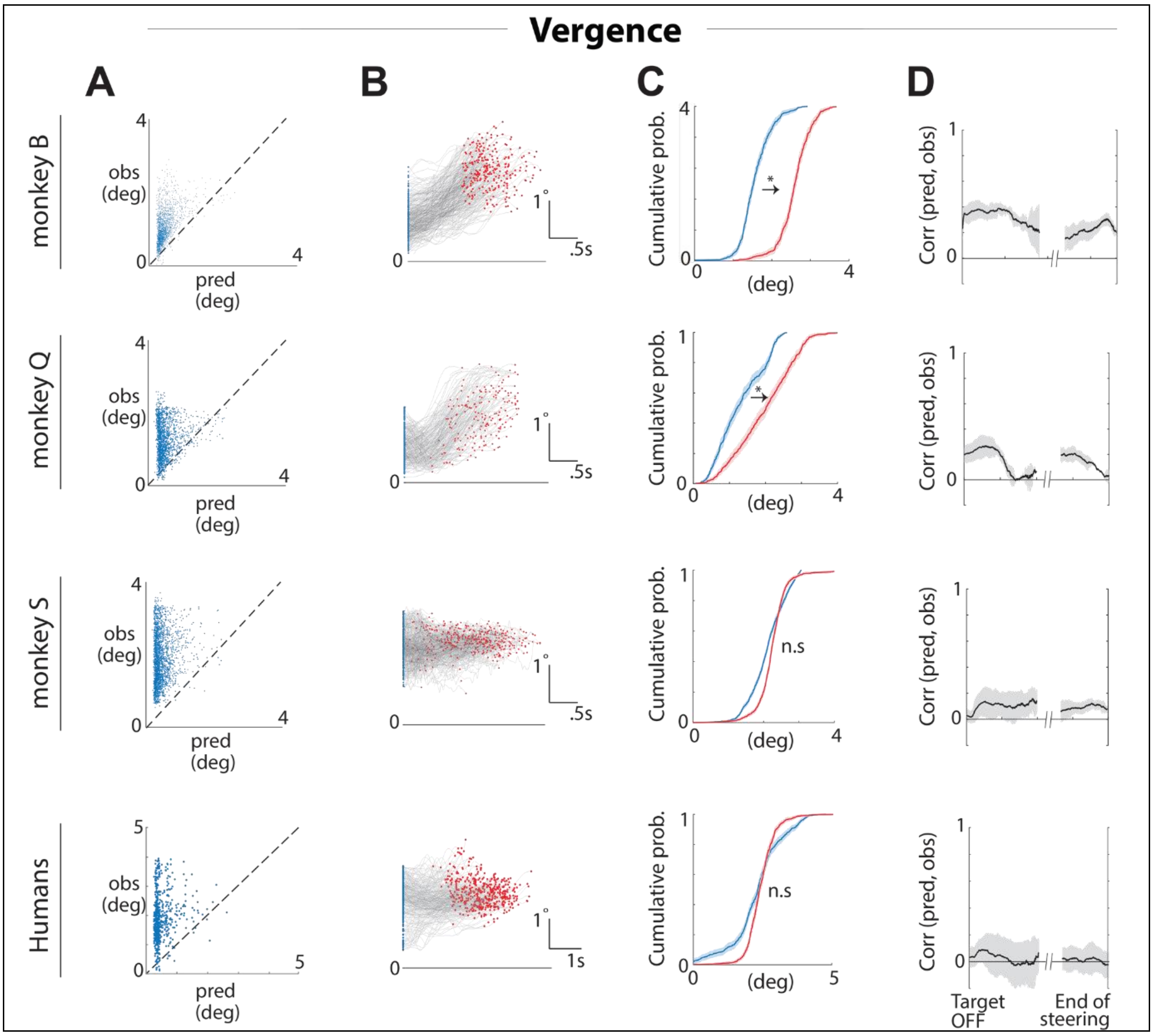

Vergence
A
B
C
D
monkey B
monkey Q
monkey S
Humans
obs (deg)
pred (deg)
Cumulative prob.
(deg)
Corr (pred, obs)
n.s
.5s
1s
Target OFF
End of steering

**Figure S7. Temporal dynamics of vergence. A.** Comparison of the predicted and true vergence in a random subset of trials sampled from individual monkeys (rows 1-3) and all human subjects (combined, last row) at the moment when the target is turned OFF. **B.** The time-course of vergence during a random subset of trials from monkeys and humans. Blue and red dots denote the times at which the target was turned OFF and the end of movement, respectively. **C.** The empirically estimated cumulative density functions of the distribution over vergence at the moment when the target was turned OFF (*blue*) and at the end of movement (*red*). Shaded regions denote standard errors obtained by bootstrapping. We used a bootstrap test with 10,000 bootstrap samples to determine whether parameters of the distributions estimated at the two time points were different. Specifically, we qualitatively expect the vergence to increase over time (*i.e.* convergence, **Fig S3**), so we tested whether the magnitude of the median was significantly greater at the end of movement. Asterisks denote significant difference ($p<0.001$) and n.s. denotes not significant according to the bootstrap test described above (*i.e.* whenever $p >= 0.001$). **D.** Time-course of the Pearson's correlation between the predicted and observed values of vergence computed by aligning trials with respect to both the time when the target was turned OFF (denoted by the y-axis on the left) as well as the end of movement (denoted by the y-axis on the right). For humans, we pooled trials from all five subjects for the analysis shown here.

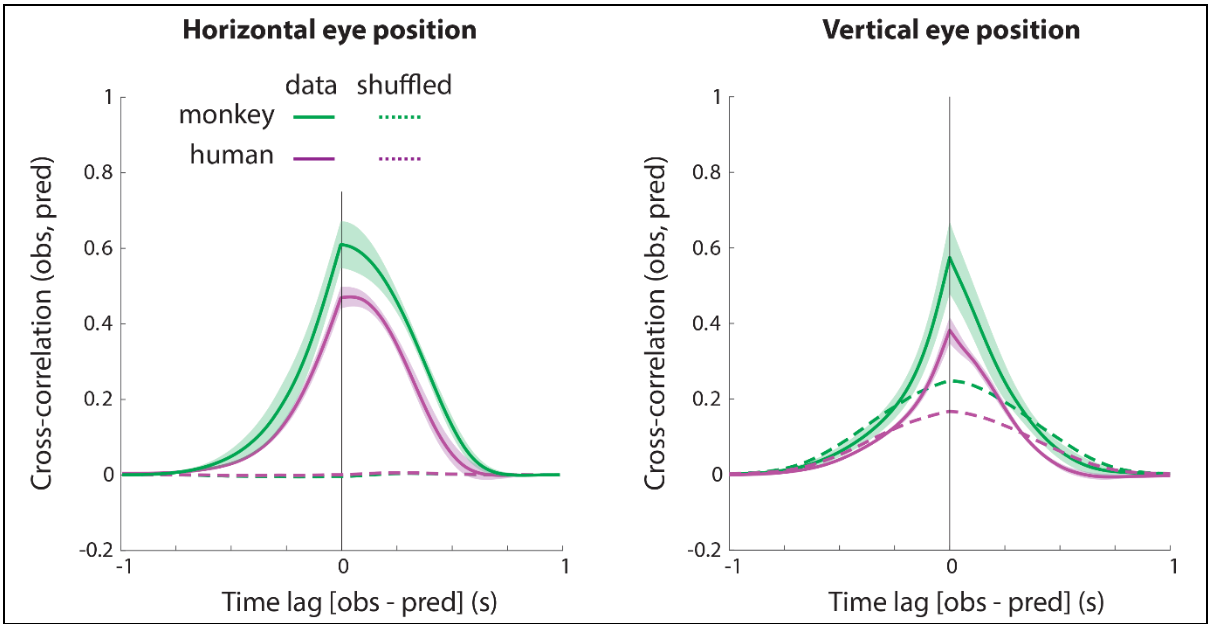


**Figure S8. Eye movements are not predictive of future target location.** *Left*: Cross-correlogram between observed and predicted horizontal eye position computed by concatenating all the trials. Error bars denote $\pm 1$ standard error in mean across subjects. A peak at positive (negative) time lag would correspond to observations lagging behind (leading) the predictions. *Right*: Cross-correlogram between observed and predicted vertical eye position. The time interval containing lower-bound of the 95% confidence interval (CI) of the peak cross-correlation did not exclude zero for both horizontal (monkeys: $\tau_{\text{peak}} = [-0.05\,, 0.15]$s ; humans: $\tau_{\text{peak}} = [-0.03\,, 0.2]$s) and vertical (monkeys: $\tau_{\text{peak}} = [-0.02\,, 0.04]$s, humans: $\tau_{\text{peak}} = [-0.03\,, 0.04]$s) components.

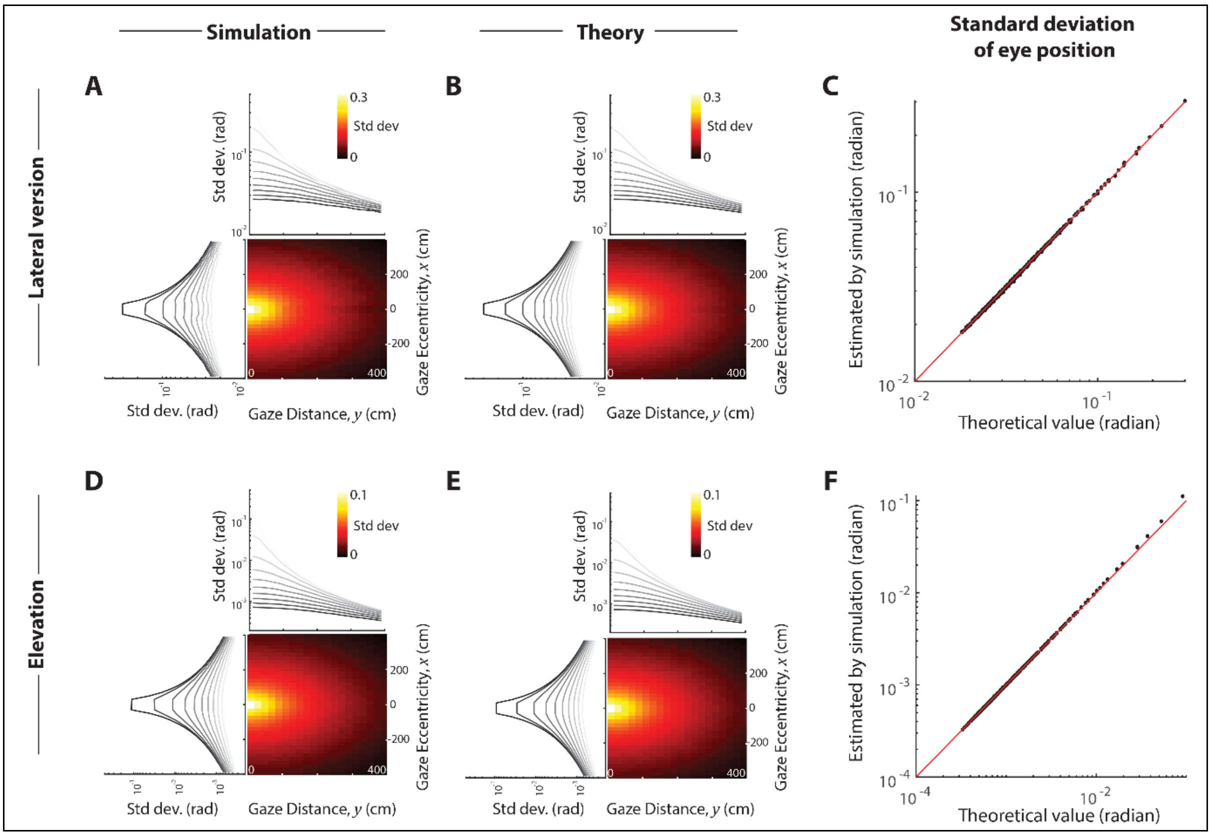


**Figure S9. Quality of theoretical approximation of variance in eye position.** We evaluated the quality of theoretical approximation of variance in eye position (**equation 3**) assuming isotropic Gaussian variance in the estimated target position $(x, y)$ on the horizontal plane ($\sigma_x = \sigma_y = 10$cm). Viewing height above the ground was held fixed ($z = 10$cm, $\sigma_z = 0$). **A.** The heatmap shows the standard deviation of horizontal eye position (*lateral version*) as a function of target position $(x, y)$ obtained by simulating **equation 1** (10,000 i.i.d. samples for each spatial bin). The plots to the top and to the left of the heatmap show dependence of standard deviation in eye position on target distance and eccentricity separately. **B.** The theoretical standard deviation in horizontal eye position calculated using **equation 4.1**. **C.** Comparison of simulated and theoretical standard deviations across all target locations. Red line has unity slope. **D-F.** Similar to **A-C**, but showing standard deviation of vertical eye position (*elevation*). We found similar correspondence between theory and simulation for other choices of parameters.

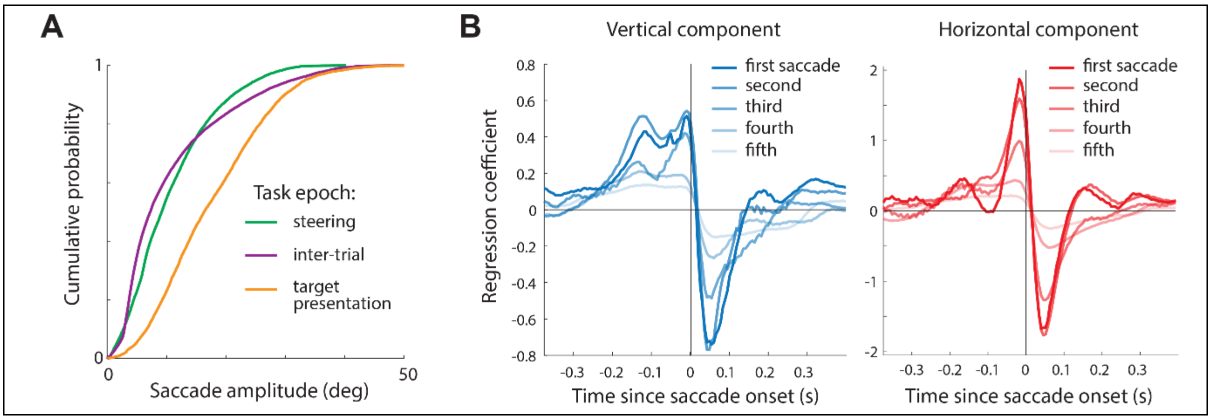


**Figure S10. Saccadic eye movements in humans. A.** Empirical cumulative distribution function of saccade amplitudes conditioned on the task epoch, averaged across all human subjects. Mean saccade amplitude ± SE: inter-trial – 10.8 ± 1.5°, target-presentation – 17.6 ± 2.2°, task phase – 10.4 ± 1.8°) **B.** The time-course of coefficients obtained by linearly regressing the amplitudes of the vertical (left) and horizontal (right) components of saccades (made while steering towards the target) against the corresponding components of the target tracking error (**Methods**). Regression was carried out separately for the first, second, third, fourth, and fifth saccades made during steering. Peak-to-peak difference in weights for vertical component: first saccade – 1.1 ± 0.3, fifth – 0.3 ± 0.1; horizontal component: first saccade – 3.6 ± 0.5, fifth – 0.5 ± 0.2.

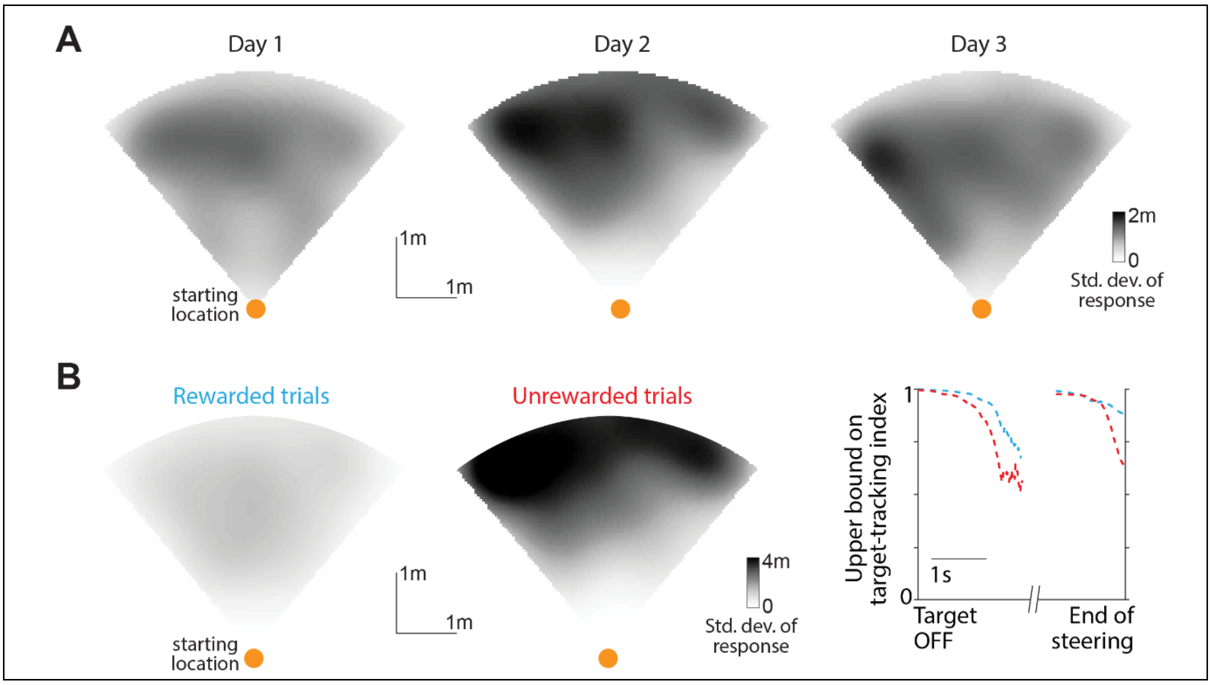


**Figure S11. Position uncertainty varied across days and across trials. A.** Aerial view of the spatial map showing the standard deviation of stopping positions as a function of target location across trials from three different sessions of one monkey. The shape of the map is due to the restricted angles and distances at which targets could appear (±40° up to 4m away). Orange dot denotes starting location. **B.** Aerial view of the spatial map showing the standard deviation of stopping positions as a function of target location across a random subset of rewarded (*left*) and unrewarded (*middle*) trials of one monkey. *Right*: Time-course of the upper bound of the target-tracking index (**equation 3**) computed separately for the two sets of trials (blue – rewarded, red - unrewarded).

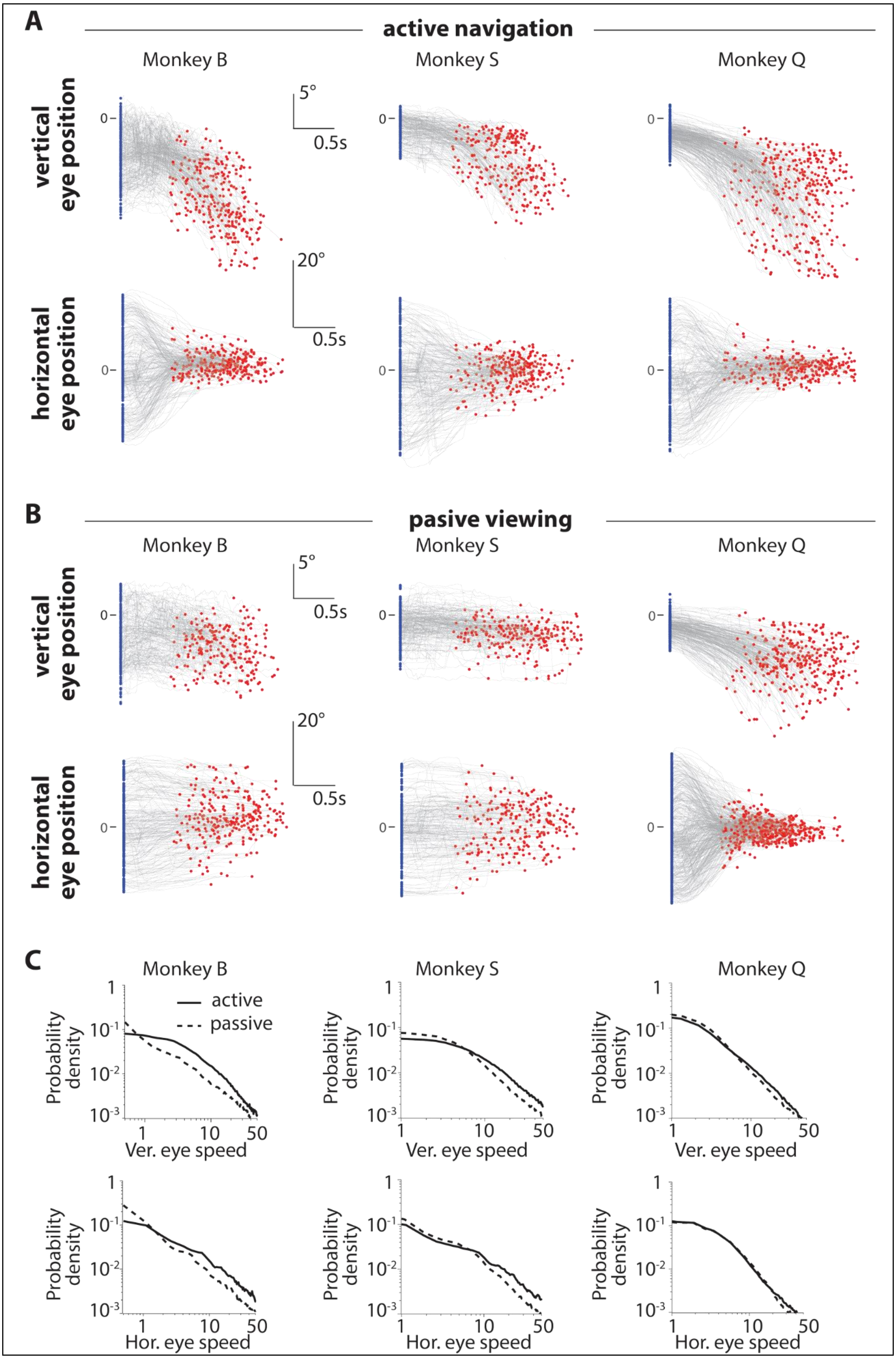

A
active navigation
Monkey B
Monkey S
Monkey Q
vertical eye position
horizontal eye position
5°
0.5s
20°
0.5s
0
B
pasive viewing
Monkey B
Monkey S
Monkey Q
vertical eye position
horizontal eye position
5°
0.5s
20°
0.5s
0
C
Monkey B
Monkey S
Monkey Q
active
passive
Probability density
1
10-1
10-2
10-3
1
10
50
Ver. eye speed
Hor. eye speed

**Figure S12. Temporal dynamics of eye position during active navigation and passive viewing (replay) blocks. A.** The time-course of elevation (top) and lateral version (bottom) during a random subset of trials from a normal block of experiment (*active navigation block*). Blue and red dots denote the times at which the target was turned OFF and the end of movement, respectively. **B.** Similar to A, but data collected during a block of trials when the movie of the visual stimulus generated during the original task was replayed to the monkeys, with joystick control withheld (*passive viewing block*). During the passive block, eye movements of monkey B and monkey S do not resemble those observed during the normal experiment. **C.** Probability distribution over vertical (top) and horizontal (bottom) eye speeds under active navigation (*solid black*) and passive viewing (*dashed black*) conditions. Note that this is a log–log plot. Median vertical eye speed: active block – $6.3 \pm 2.8\ °/s$ ; passive block – $3.9\ \pm 2.7\ °/s$. Median horizontal eye speed: active block – $5.7 \pm 2.3\ °/s$ ; passive block – $2.8\ \pm 1.3\ °/s$.

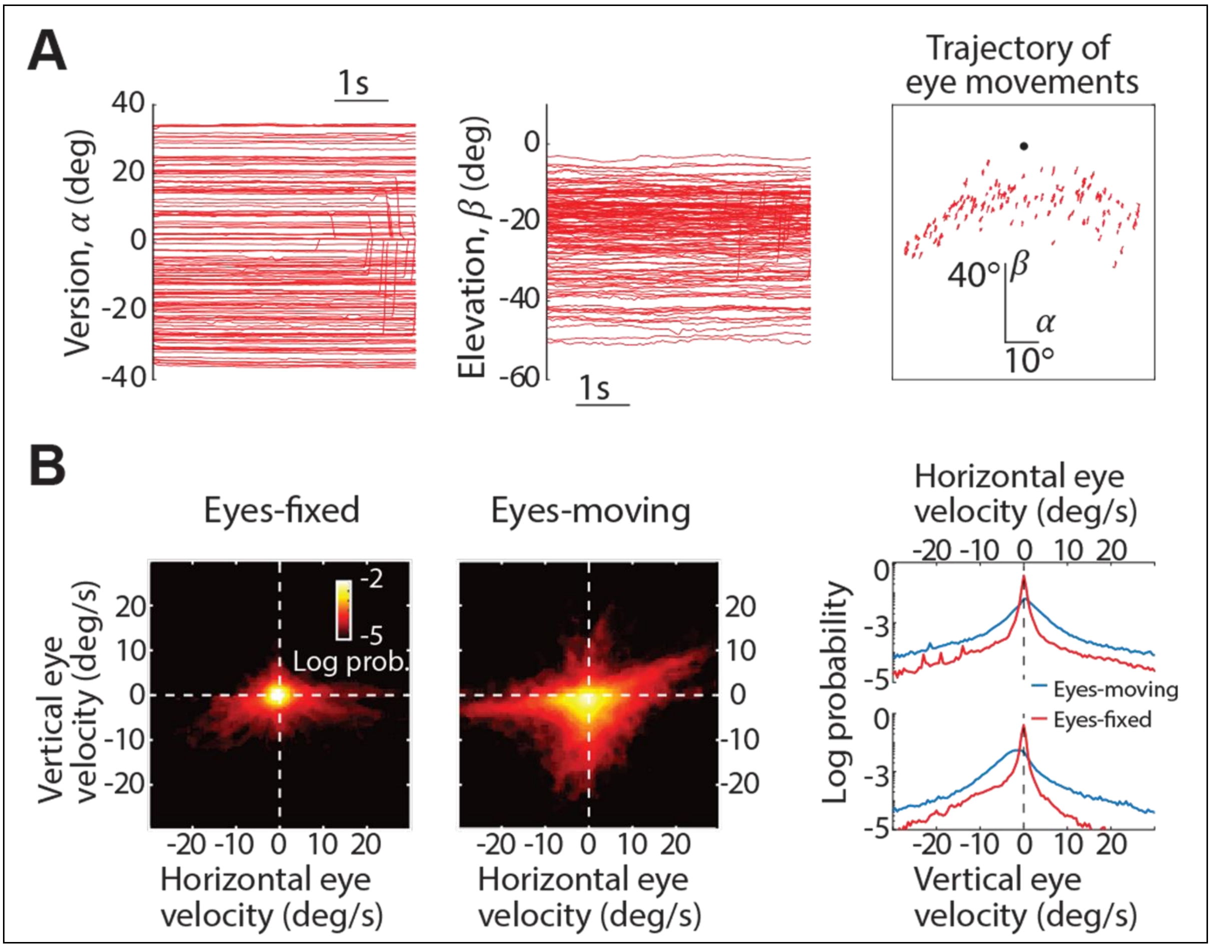


**Figure S13. Eye movements during the fixation task. A.** Time-course of lateral version, $\alpha$ (left) and elevation, $\beta$ (middle) of one human subject during a subset of trials in the 'Eyes-fixed' condition. The rightmost panel shows the trajectory (time-course) of eye movements in the $\alpha$-$\beta$ space during the same set of trials. Black dot denotes the origin $(\alpha, \beta) = (0,0)$. For this subject, the mean (± std) temporal variability (quantified as standard deviation $\sigma$ of eye position across time, see **Methods**) of eye movements during the 'Eyes-fixed' condition was $0.5° \pm 0.3°$. See **Fig 5A** for summary data of all subjects. **B.** *Left*: The joint probability density of the distribution over the subjects' horizontal and vertical eye velocity, averaged across all human subjects, separately for trials from 'Eyes-fixed' and 'Eyes-moving' conditions. *Right*: Marginal distributions over horizontal and vertical eye velocity under the same two conditions. Note the much larger concentration of values around zero velocity during the 'Eyes-fixed' condition implying that subjects did not move their eyes during this condition.
